# Tethering-facilitated DNA ‘opening’ and complementary roles of β-hairpin motifs in the Rad4/XPC DNA damage sensor protein

**DOI:** 10.1101/2020.09.28.313049

**Authors:** Debamita Paul, Hong Mu, Amirrasoul Tavakoli, Qing Dai, Xuejing Chen, Sagnik Chakraborty, Chuan He, Anjum Ansari, Suse Broyde, Jung-Hyun Min

## Abstract

XPC/Rad4 initiates eukaryotic nucleotide excision repair on structurally diverse helix-destabilizing/distorting DNA lesions by selectively ‘opening’ these sites while rapidly diffusing along undamaged DNA. Previous structural studies showed that Rad4, when tethered to DNA, could also open undamaged DNA, suggesting a ‘kinetic gating’ mechanism whereby lesion discrimination relied on efficient opening versus diffusion. However, solution studies in support of such a mechanism were lacking and how ‘opening’ is brought about remained unclear. Here, we present crystal structures and fluorescence-based conformational analyses on tethered complexes, showing that Rad4 can indeed ‘open’ undamaged DNA in solution and that such ‘opening’ can largely occur without one or the other of the β-hairpin motifs in the BHD2 or BHD3 domains. Notably, the Rad4-bound ‘open’ DNA adopts multiple conformations in solution notwithstanding the DNA’s original structure or the β-hairpins. Molecular dynamics simulations reveal compensatory roles of the β-hairpins, which may render robustness in dealing with and opening diverse lesions. Our study showcases how fluorescence-based studies can be used to obtain information complementary to ensemble structural studies. The tethering-facilitated DNA ‘opening’ of undamaged sites and the dynamic nature of ‘open’ DNA may shed light on how the protein functions within and beyond NER in cells.

## INTRODUCTION

Nucleotide excision repair (NER) is an evolutionarily conserved DNA repair pathway that protects the genome integrity against environmental mutagens including UV light and a wide variety of man-made and natural chemicals (1,2). The key to the versatility of NER lies in its unique lesion recognition and initiation mechanism involving the XPC-RAD23B-CETN2 complex (hereafter XPC). Using thermal energy alone, XPC surveys DNA within chromatin and locates diverse target lesions by specifically binding to them (3–8). DNA lesions that XPC recognizes for NER include various intra-strand crosslinks and helix destabilizing/distorting adducts, which are formed by ultraviolet (UV) light, air and water pollutants, and toxins (9). Once bound to a lesion, XPC in turn recruits the 10-subunit general transcription factor II H complex (TFIIH), which then verifies the presence of a bulky lesion and recruits other subsequent factors (10–13); eventually the lesion-containing single-stranded DNA is excised by XPF-ERCC1 and XPG endonucleases and the DNA is restored by repair synthesis and nick sealing by DNA polymerases and DNA ligases, respectively (9,14). Genetic mutations in XPC and other NER factors underlie diseases such as the xeroderma pigmentosum (XP) cancer predisposition syndrome, marked by extreme sun sensitivity and >1000-fold higher risk of sunlight-induced skin cancers (15–17). Recent genome-wide mapping of DNA damage and cancer mutations also showed that inefficient NER is a major contributor to mutation hotspots in sporadic skin and lung cancers (18–23).

A key to the function of XPC as a versatile damage sensor lies in its ability to search and scan along the DNA and then selectively ‘open’ a DNA duplex where there is damage. The previous crystal structures of the yeast XPC ortholog, Rad4, bound to UV-induced 6-4 photoproduct (6-4PP) or mismatched DNA showed that Rad4 (and by analogy XPC) recognizes the damaged DNA site by inserting a β-hairpin from its β-hairpin domain 3 (BHD3) into the DNA duplex and flipping out damage-containing nucleotide pairs to form an “open” conformation (24,25). In this conformation, Rad4 interacts exclusively with the two nucleotides on the undamaged strand without making specific contacts to the two damaged residues. This indirect recognition thus can allow Rad4/XPC to bind to DNA lesions with varied structures (24,26). Intriguingly, we also observed that this ‘open’ conformation could be formed on undamaged DNA in crystals when Rad4 was site-specifically tethered to DNA to minimize heterogeneity in the binding registers, inherent in nonspecific complexes (27). The tethering did not induce any structural distortions in the protein-DNA contacts and was also site-specific. However, together with subsequent biophysical kinetics studies, our results suggested that Rad4/XPC may rely on a ‘kinetic gating’ mechanism (27–29). In this mechanism, damaged DNA may be selectively ‘opened’ over undamaged DNA by a freely diffusing protein, not necessarily because the protein is inherently unable to ‘open’ undamaged DNA but because the free energy barrier for opening is high and the protein does not linger long enough at an undamaged site to efficiently ‘open’ that site before diffusing away (27–29). On the other hand, damaged DNA, since already destabilized, has a lower free energy barrier for opening, thus increasing the probability that the protein can ‘open’ that site and not diffuse away. Such a kinetically controlled, indirect readout mechanism presents a paradigm on how specific DNA recognition can be brought about, which enables rapid, yet reliable recognition of diverse lesions without wasteful interrogation of the predominantly nonspecific undamaged background as in genomic DNA.

Despite these studies, however, observation of the ‘opening’ of undamaged DNA by Rad4 has been limited to the crystal structure and evidence for such an ‘open’ complex conformation in solution has been lacking. It has also been unclear how the duplex unwinding and ‘opening’ is brought about by Rad4/XPC. Here, we examined the ‘opening’ mechanism with a special focus on the roles of the β-hairpins from BHD2 and BHD3 that engage DNA at the ‘open’ site, using various complementary approaches that include crystallography, fluorescence studies, and molecular dynamics (MD) simulations. The crystal structures of Rad4 mutants lacking the β-hairpins in either BHD2 or BHD3 (Δβ-hairpin2 and Δβ-hairpin3, respectively) showed that the mutants, when tethered, could also open undamaged DNA in a manner similar to wild-type (WT) Rad4. To seek evidence for such ‘open’ complex conformations in solution, we then turned to a fluorescence lifetime (FLT)-based conformational analysis approach with a unique set of fluorescence resonance energy transfer (FRET) pair (cytosine analogs tC^o^ and tC_nitro_) incorporated in the DNA duplexes; these FRET probes have proven to be exquisitely sensitive to small changes in DNA helical conformations (29–34). These FLT-FRET studies revealed similar ‘open’ conformations with undamaged DNA (CCC/GGG) tethered to WT Rad4 as with damaged DNA (3-bp CCC/CCC mismatch) noncovalently bound to WT Rad4. The ‘open’ conformations of the DNA tethered to mutant Rad4 were also similar to those of the WT complexes, further corroborating the crystal structures. Surprisingly, the Rad4-bound ‘open’ DNA sampled multiple conformations with similar population distributions regardless of whether the DNA was damaged or not or the absence/presence of the β-hairpins. Subsequent MD simulations showed that β-hairpin2 facilitates further DNA untwisting in the absence of β-hairpin3, while β-hairpin3 promotes additional partner base extrusion in the absence of β-hairpin2, revealing the compensatory roles of the β-hairpins and suggesting that DNA ‘opening’ need not be reached via the same trajectory.

To the best of our knowledge, this study represents the first 3-D structural analyses of any mutant Rad4/XPC and also showcases how crystallographic structures and solution fluorescence experiments can provide valuable complementary information concerning structural mechanisms. The dynamically fluctuating ‘open’ DNA conformations may be important for the downstream processes of NER and the complementary roles of multiple β-hairpins may render robustness to the mechanism for Rad4/XPC in dealing with diverse lesions. Our study fills in key missing pieces in the kinetic gating mechanism picture and provides an important step towards structural understanding of how Rad4/XPC brings about the indirect recognition of diverse NER lesions.

## MATERIALS AND METHODS

### Preparation of Rad4–Rad23 complexes

The intact (‘WT’) and the Δβ-hairpin3 Rad4-Rad23 complex constructs are as published previously (24,27–29). Rad4 in the WT construct spanned residues 101–632 and contained all four domains involved in DNA binding, exhibiting the same DNA-binding characteristics as the full-length Rad4-Rad23 complex (24). The Δβ-hairpin3 mutant complex (construct name: <137>) lacked the β-hairpin in the BHD3 domain (residues 599-605) of Rad4 whereas Δβ-hairpin2 (<130>) lacked the β-hairpin in BHD2 (residues 515-527) in the context of the WT construct. All complexes contained the same Rad23 construct comprising the UBL, Rad4-binding and UBA2 domains, as before (24,27–29). For preparation of the crosslinked Rad4-Rad23 complexes with DNA, the ‘WT’ (<SC32>), Δβ-hairpin3 (<SC41b>) and Δβ-hairpin2 (<SC40>) constructs also harbored V131C/C132S mutations in Rad4 to allow site-specific disulfide crosslinking with the dG*-containing DNA as done before (27).

The Rad4–Rad23 complexes were co-expressed and purified from baculovirus-infected insect cells as before (24). Briefly, the Hi5 insect cells co-expressing Rad4 and Rad23 were harvested two days after infection. After lysis, the proteins were purified using immobilized metal affinity chromatography (Ni-NTA agarose, MCLAB) and then anion-exchange chromatography (Source Q, GE healthcare). The complexes were then subjected to thrombin digestion at 4 °C overnight, followed by cation exchange (Source S, GE healthcare) and size-exclusion (Superdex200, GE healthcare) chromatography. The purified proteins were concentrated by ultracentrifugation (Amicon Ultra-15, Millipore) to ~15-20 mg/ml and stored in 5 mM bis-tris propane–HCl (BTP-HCl), 800 mM sodium chloride (NaCl) and 5 mM dithiothreitol (DTT), pH 6.8. For competitive EMSA and fluorescence studies, the protein complexes were purified without thrombin digestion, thus retaining the UBL domain of Rad23 and an N-terminal histidine-tag on Rad4 as previously described (24,27).

### Synthesis of oligonucleotides and preparation of duplex DNA

All unmodified oligonucleotides were purchased from IDT as HPLC-purified. Oligonucleotides modified with tC^o^ and tC_nitro_ were purchased from Biosynthesis as HPLC-purified. In general, duplex DNA samples were prepared by mixing two complementary oligonucleotides in 1:1 ratio and annealing by slow-cooling in water. The oligonucleotides containing a disulfide-modified guanine (dG* in the top strand) used for crystallization or fluorescence measurements were prepared by incorporating the 2-F-dI-CE phosphoramidite (Glen Research) at the desired position during solid-phase synthesis. The conversion and deprotection of 2-F-dI were performed according to Technical Bulletin provided by Glen Research (https://www.glenresearch.com/media/productattach/import/tbn/TB_2-F-dI.pdf). Briefly, 2-F-dI-containing oligonucleotides were treated with cystamine (prepared freshly from cystamine hydrochloride and sodium hydroxide) to tether the disulfide group, then deprotected with 1,8-diazabicycloundec-7-ene. The resulting dG*-containing oligonucleotides were purified with HPLC and verified by MALDI-MS. The DNA duplexes containing dG* were prepared by annealing the top and bottom strands in 1:1 (fluorescence lifetime) or 1:1.1 (crystallization) ratios in water.

### Site-specific disulfide tethering

First, DTT from the Rad4-Rad23 samples was removed by buffer exchange using a desalting column (Zeba Spin Desalting Column, 40,000 Da molecular weight cut-off, Thermo Scientific) to final buffer containing 5 mM BTP-HCl and 800 mM NaCl, pH 6.8. The protein complexes were subsequently mixed with the dG*-containing DNA duplex in 1:1 molar ratio in crosslinking buffer (5 mM BTP-HCl, 100 mM NaCl and 10% glycerol, pH 6.8) and were incubated at 4 °C overnight. The reaction progress was determined by SDS–PAGE under non-reducing conditions after treating the samples with 0.1 mM S-methylmethanethiosulfonate (Sigma) to quench the reaction. The crosslinking yield was typically ~50–60%. The complexes were then purified by anion-exchange chromatography (Mono Q, GE healthcare) over a 0–2 M NaCl gradient in 5 mM BTP-HCl and 10% glycerol, pH 6.8, in which the buffers were degassed continually by nitrogen purging. Crosslinked protein-DNA complexes eluted at 400–480 mM NaCl, well separated from free protein and free DNA, and the purified samples were further concentrated by ultrafiltration (Amicon, Millipore).

### Crystallization and x-ray diffraction data collection

All crystals were grown by the hanging-drop vapor diffusion method at 4 °C in which 1.5 μl of protein-DNA complex was mixed with 1.5 μl of crystallization buffer (50 mM BTP-HCl, 150 mM NaCl and 10–15% isopropanol, pH 6.8) and sealed over 1 ml of crystallization buffer. Showers of needle-like small crystals (10-20 μm) appeared within a few days. To obtain larger crystals, micro-seeding was carried out: hanging drops with 1:1 mix of the protein-DNA sample and crystallization buffer (50 mM BTP-HCl, 180 mM NaCl and 5-15% isopropanol, pH 6.8) were pre-equilibrated for 1 h, then seeded by passing the tip of a cat whisker dipped into a fresh seed-stock solution made up of small microcrystals across the drop. The crystals grew to maximum size of ~30-50 μm in 10-12 days. The crystals were harvested in a harvest buffer (50 mM BTP-HCl, 180 mM NaCl, 12% isopropanol) and submerged for few seconds in a cryoprotectant buffer (50 mM BTP-HCl, 180 mM NaCl, 12% isopropanol, 5% glycerol and 20% 2-methyl-2,4-pentanediol) before flash-frozen in liquid nitrogen. Diffraction data were collected in LS-CAT (21-ID-F) beamline at 103 K at the Advanced Photon Source and were processed with the HKL2000. The data collection statistics are summarized in **Table S1**.

### Structure determination and refinement

The structure of the mutant Rad4–Rad23–DNA complexes were determined by molecular replacement method using the previous structure of WT Rad4 crosslinked with CCC/GGG only using the protein portion first (chains A and X in 4YIR (PDB ID)) as the search model using MOLREP (CCP4). The DNA was searched separately after fixing the protein in the unit cell, using Phaser (CCP4) (35). Several rounds of model buildings were performed using WinCoot (36) followed by refinements with Phenix (37).

The refinement statistics are summarized in **Table S1**. The final structure of Δβ-hairpin3-DNA (PDB ID: 6UBF) contains residues 126–516, 525–596 and 607-632 of Rad4 and 256–308 of Rad23. The DNA residues with missing densities are W16-17 in top strand and Y9-11 in bottom strand. The final structure of Δβ-hairpin2-DNA (PDB ID: 6UIN) contains residues 130–513, 528–599 and 606-632 of Rad4 and 255-308 of Rad23. The DNA residues with missing densities are W15-17 in top strand and Y8-11 in bottom strand. All structural figures were made using PyMOL Molecular Graphics System, version 2.1.1 (Schrodinger, LLC).

### Apparent binding affinities (K_d,app_) determined by competition electrophoretic mobility shift assays (EMSA)

The specified Rad4-Rad23 complexes (0-300 nM) were mixed with 5 nM ^32^P-labelled DNA substrate in the presence of 1000 nM cold, undamaged competitor DNA (CH7_NX) in a binding assay buffer (5 mM BTP-HCl, 75 mM NaCl, 5 mM DTT, 5% glycerol, 0.74 mM 3-[(3-cholamidopropyl)dimethylammonio]-1-propanesulfonate (CHAPS), 500 µg/ml bovine serum albumin (BSA), pH 6.8). Mixed samples were incubated at room temperature for 20 min before being separated on 4.8% non-denaturing polyacrylamide gels (10-15 min at 150 V). The gels were quantitated by autoradiography using Typhoon FLA 9000 imaging scanner (GE healthcare) and Image Lab software (Version 5.2.1 build 11, 2014; Bio-Rad). The apparent dissociation constants (K_d,app_) were calculated as previously described using curve fitting by nonlinear regression using Origin software (OriginPro 9.6.0.172, Academic) (25,27,29,33).

### Time-resolved fluorescence spectroscopy for fluorescence lifetime (FLT) measurements

DNA duplexes labeled with both tC^o^ and tC_nitro_ or tC^o^ alone were prepared as described above. The DNA and protein-DNA samples were prepared at 5 μM in phosphate-buffered saline (PBS: 10 mm Na_2_HPO_4_, 2 mM KH_2_PO_4_, 137 mM NaCl, 2.7 mM KCl pH 7.4) with 1 mM DTT.

Sample volume for each FLT measurement was 45 μl. Fluorescence decay curves for the FRET donor tC^o^ (in the absence and presence of the FRET acceptor tC_nitro_, which in itself is nonfluorescent) were measured with a DeltaFlex fluorescence lifetime instrument (HORIBA) equipped with a Ti-sapphire laser source as an excitation source (Mai Tai HP, Spectra-Physics). The beam for tC^o^ excitation was produced by frequency doubling of the fundamental beam (730 nm) and pulse-picking at 4 MHz, which was then passed through a monochromator set at 365 nm (band pass 10 nm). The fluorescence signal emitted at 470 nm (band pass 10 nm) was collected by a Picosecond Photon Detection module (PPD-850, Horiba) using time-correlated single-photon counting (TCSPC) electronics. Fluorescence decay curves were recorded on a 100 ns timescale, resolved into 4,096 channels, to a total of 10,000 counts in the peak channel. All details are in SI Methods.

### Fluorescence lifetime analyses using maximum entropy method (MEM) and Gaussian fitting

The decay traces were analyzed by the maximum entropy method (MEM) using MemExp software (38,39), as done previously (40). The MEM analysis yielded a distribution of donor lifetimes with each lifetime component (*τ_D,i_* for donor-only samples and *τ_DA,i_* for donor-acceptor labeled samples) having a corresponding amplitude *α_i_*. The reproducibility of the distributions obtained from the MEM analyses are illustrated from 2-3 independent lifetime measurements on each sample in **Figure S6**. The lifetime distributions from the MEM analyses were subsequently fitted as a sum of Gaussians (**Figure S6**). The donor-only samples exhibited a single peak and *τ_D_*, the characteristic lifetime of the donor-only sample, was obtained from the peak position of the Gaussian-fitted distribution. The average FRET efficiency for each sample was computed as 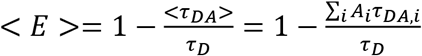, where 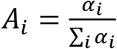 is the normalized amplitude corresponding to each lifetime component (*τ_DA,i_*). Each Gaussian component in the *τ_DA_* distribution was used to calculate the average lifetime and FRET efficiency representing that component, and the area under the Gaussian curve was taken as a measure of the fractional population of that component. The results are summarized in **Table S2.** Errors are indicated with standard deviations (s.d.) from 2-3 independent sets of measurements.

### Molecular dynamics simulations

Δβ-hairpin2 and Δβ-hairpin3 were modeled with truncation in the respective β-hairpin tips to match the mutant constructs used in the experimental studies. The initial models of the proteins docked to the CCC/GGG DNA were prepared similarly as before where the BHD2 and BHD3 domains are close but not yet bound to the DNA duplex (25,41). During the 2 μs MD simulations, stable BHD2 conformations were achieved at ~1 µs (**Figure S9**). Hence the 1–2 µs trajectories were taken as the initial binding states (IBS) for each complex and analyzed for structural, energetic and dynamic properties. The untwisting, bend angle, bend direction pseudo-dihedral angle around the potential ‘open’ site (base pair step W16-17, **Figure 2A**) and the extent of BHD2 binding into the DNA minor groove were measured. The untwisting of 6-mer (base pair steps W14 and W19) was monitored by the Untwist angle defined as Untwist = Twist _initial_ –Twist. Twist _initial_ is the ensemble average twist angle of the lesion-containing 6-mer during the first 1 ns of production MD during which significant untwisting was not observed (**Figure S10**); this ensemble represents the state of the lesion-containing sequence before the engagement of Rad4, especially BHD2. Positive values indicate further untwisting and negative values indicate further twisting. Bend angles and bend direction dihedral angles are defined in **Figure S10**. The BHD2 occupied AlphaSpace of the DNA minor groove was measured as previously described (25,41) to reflect the extent of BHD2 binding. Base extrusion for the potential flipping nucleotides (dC’s in W16-17, **Figure 2A**) was measured using a pseudo-dihedral angle (**Figure S11**). The van der Waals interaction energy and hydrogen bonds for BHD2/3 residues that interact with the DNA were also computed to identify essential residues in BHD2/3 functions. All details of force field, MD simulations and structural analyses are given in SI Methods, and details of additional sampling are given in SI Results.

## RESULTS

### Rad4 mutants lacking β-hairpins from BHD2 or BHD3 bind more weakly to DNA compared with WT with reduced specificities for mismatches

Mismatched bubbles such as CCC/CCC can be recognized by Rad4/XPC *in vitro* in a manner similar to UV-induced photolesions such as 6-4 photoproducts repaired by NER, and thus have served as useful models for investigating Rad4/XPC interactions with damaged DNA (24,27,42). To examine the roles of the β-hairpins in influencing the protein-induced DNA opening, we have prepared Rad4-Rad23 mutant complexes lacking the tip of the β-hairpins from either the BHD2 or the BHD3 domains of Rad4 (Δβ-hairpin2 and Δβ-hairpin3) and first tested their binding to DNA in a competitive electrophoretic mobility shift assay (EMSA) as previously described (**Figure 1**) (24,25,27,29,34). The apparent dissociation constant (K_s,app_) for the intact Rad4 (‘WT’) binding to a specific DNA substrate containing a 3-bp mismatch CCC/CCC was ~60 nM, which was ~6.2 fold lower than that to a corresponding matched DNA substrate (CCC/GGG) for which the apparent dissociation constant (K_ns,app_) was ~370 nM. On the other hand, the Δβ-hairpin3 mutant showed only a ~2.5-fold lower affinity to the matched DNA than to the mismatched DNA (K_s,app_ ~200 nM versus K_ns,app_ ~500 nM), and a ~1.4-fold lower affinity than that of WT for the matched DNA. These results for WT and the Δβ-hairpin3 mutants were consistent with our previous reports using the same assay (24,27). Interestingly, Δβ-hairpin2 also showed a ~2.5-fold lower affinity for the matched DNA versus mismatched DNA (K_s,app_ ~266 nM versus K_ns,app_ ~650 nM), closely mimicking the characteristics of Δβ-hairpin3. Thus, the mutants exhibit moderately weakened DNA binding to matched DNA in comparison with WT with a significant loss in mismatch (lesion)-binding specificity; the Δβ-hairpin2 and Δβ-hairpin3 mutants were affected to similar degrees compared with WT.

**Figure 1.**
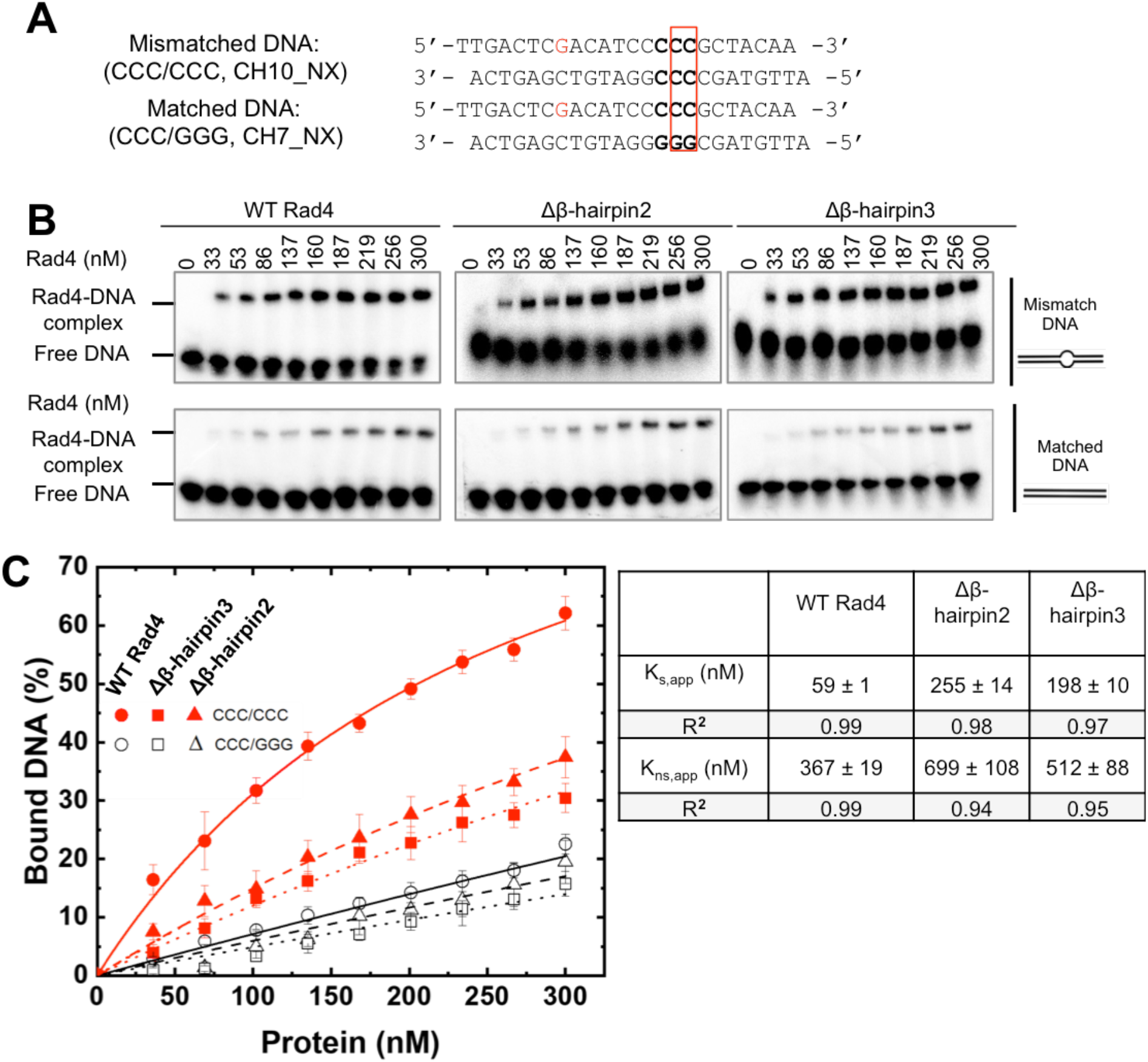
Characterization of DNA binding affinities by Rad4-Rad23 complexes. **(A)** Sequences of DNA constructs used for the competitive EMSA. The position of the two flipped-out nucleotide pairs in the ‘open’ conformation is indicated with a red box. **(B)** Typical gel images showing the various Rad4 constructs binding to the mismatched CCC/CCC (top) and matched CCC/GGG DNA (bottom). **(C)** Quantification of the percent bound DNA fractions in (B) versus protein concentrations. The symbols and error bars indicate the means and ranges as calculated by ± sample standard deviation, respectively, from triplicate gel shift experiments. Filled red symbols indicate mismatched DNA and empty black ones indicate matched DNA; circle, square and triangle indicate WT, Δβ-hairpin2 and 3, respectively. Lines indicate the fit curves of the data points. Apparent dissociation constants for specific binding to mismatched DNA (K_s,app_) and nonspecific binding to matched DNA (K_ns,app_) are also shown.

### Crystal structures of Δβ-hairpin2 and Δβ-hairpin3 tethered to CCC/GGG matched DNA show conformations mimicking the ‘open’ structure with WT Rad4

To examine the structural impacts of these mutations, we next sought to examine the crystal structures of these mutants bound to DNA. As shown by competitive EMSA described above, the mutants have significantly lower lesion-binding specificity than WT Rad4. Thus, under freely diffusing conditions, the mutants are prone to bind to DNA in multiple different registers, which hampers crystal diffraction and structure determination. Hence, we decided to use site-specific disulfide tethering to limit the binding register of the protein on the DNA as previously done. Diffracting crystals could be obtained using this technique between WT Rad4 and nonspecific, matched DNA containing the CCC/GGG sequence (27). Notably, as mentioned in the Introduction, the structure of the tethered complex showed that WT Rad4 could also ‘open’ matched DNA to form a structure similar to that seen with untethered Rad4 naturally bound to specific substrates such as the 6-4PP UV lesion or 3-bp mismatched DNA (24,25). Therefore, to make straightforward comparisons between the mutants and WT, we used the same matched CCC/GGG DNA construct together with the tethering strategy to obtain the mutant crystal structures with DNA. The tethering site is located in the N-terminus of Rad4’s TGD domain (Cys131), away from the BHD2/3 domains where the DNA duplex ‘opening’ occurs. The tethered complexes were purified to homogeneity and yielded diffracting crystals (**Figures S1 & 2, Table S1**).

Despite the differences in the space groups (*P4_1_2_1_2* for Δβ-hairpin3 and *P6_5_* for Δβ-hairpin2) and crystal packing (**Figure S2**), these mutant-DNA structures both showed protein and DNA conformations that superpose well with the previously reported ‘open’ conformation of the Rad4-Rad23-DNA complexes (**Figures 2 & 3**) (24,25,27). The root mean squared deviations (RMSD) from the WT Rad4-DNA complex structure were 0.95 Å in the Δβ-hairpin3-DNA complex and 0.90 Å for the Δβ-hairpin2-DNA complex, calculated for common 8840 atoms in the protein and DNA. Additionally, the distance between the crosslinked residues, Cα of C131 in Rad4 and C2 of G*8 in the top strand of the DNA was 8.6 Å for Δβ-hairpin3 and 8.7 Å for the Δβ-hairpin2 structure, close to the equivalent distance (9.0 Å) in the WT Rad4-6-4PP structure. These results demonstrate that the crosslinking did not introduce structural distortions in the protein-DNA contacts and was also site-specific: such a DNA-binding mode would not have been possible if tethering had occurred with any one of the other cysteines in Rad4 exposed to the solvent (C276, C354, C355, C463, C466, C509 and C572). These observations are also consistent with those observed before with WT Rad4 (27).

**Figure 2.**
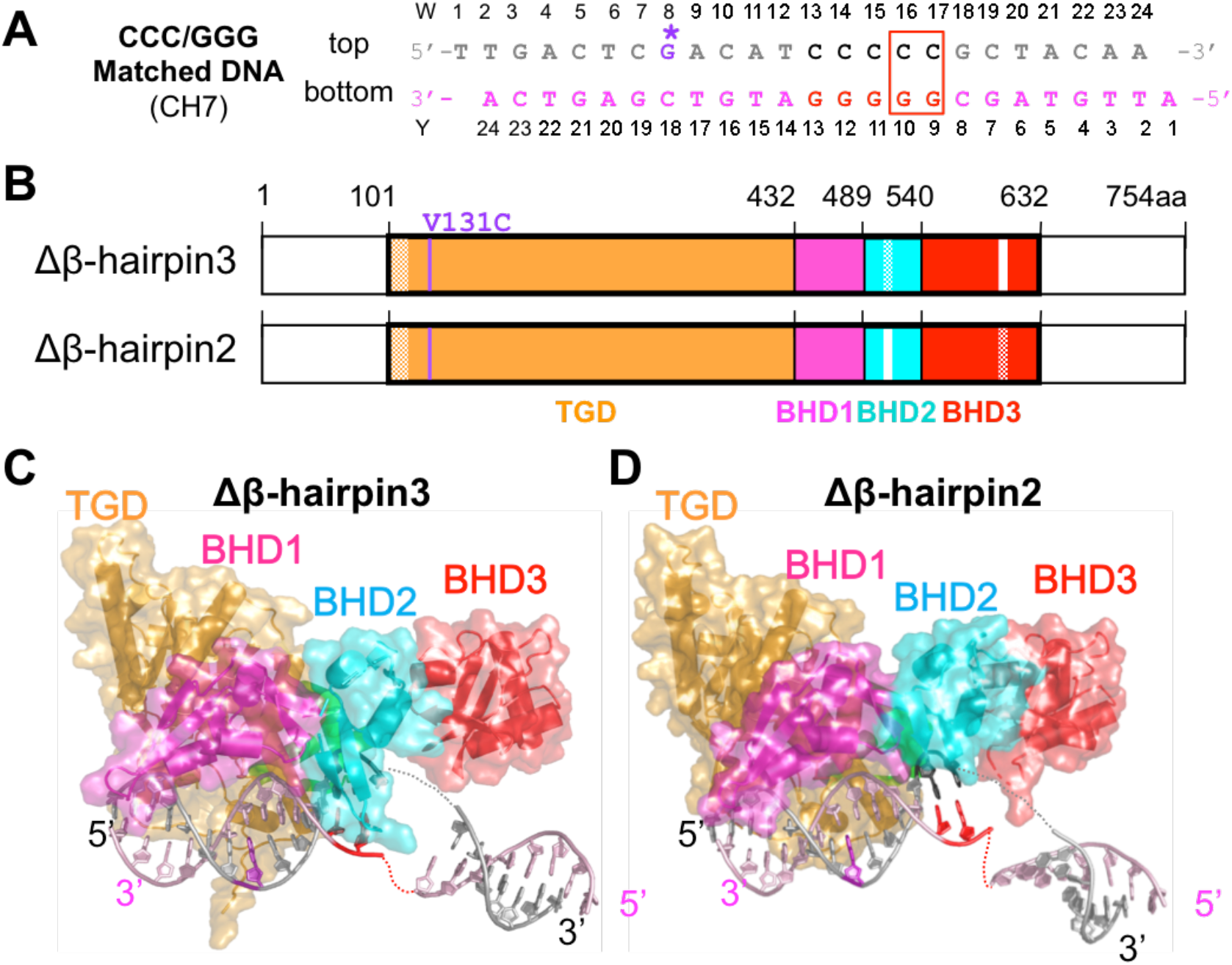
Overall structures of the β-hairpin deletion mutants of Rad4–Rad23 complexes tethered to CCC/GGG matched DNA. **(A)** The crosslinkable DNA used for crystallization. The ‘top’ strand (W) is colored in grey and the ‘bottom’ strand (Y) in light pink. The stretch of C/G’s within the sequence is in black and red. The disulfide modified G* is in purple with its chemical structure indicated in the inset. The position of the two flipped-out nucleotide pairs in the ‘open’ conformation is indicated with a red box. **(B)** The domain arrangements and boundaries of Rad4 used in this study. The transglutaminase domain (TGD) of Rad4 is indicated in orange, β-hairpin domain 1 (BHD1) magenta, BHD2 cyan and BHD3 red. Deleted β-hairpin3 (residues 599-605) in Δβ-hairpin3 and β-hairpin2 (residues 515-527) in Δβ-hairpin2 are indicated in white. Disordered regions in crystals are checkered. The V131C point mutation introduced for disulfide crosslinking is in purple. Rad23 construct was the same as in ref. (24). **(C, D)** The overall crystal structures of Δβ-hairpin3–DNA (PDB ID: 6UBF) and Δβ-hairpin2–DNA (PDB ID: 6UIN). The color codes are the same as in (A) and (B).

A salient difference between the mutant and WT structures was in the electron density around the DNA nucleotide pairs at the ‘open’ site (W16-17, **Figure 2A**). So far, all the WT Rad4 crystal structures solved (either with specific DNA substrates (e.g., 6-4PP) without tethering or with the matched CCC/GGG DNA with tethering) showed the same ‘open’ conformation (24,25,27). In such ‘open’ structures, the electron densities for the two nucleotides on the top strand (W16-17) were clearly visible as they were fully flipped out of the DNA duplex and snugly accommodated between BHD2 and BHD3, whereas the densities of the two on the bottom strand (Y9-10) were not visible as they were disordered (e.g., red and green in **Figure 3**). In comparison, the mutant-DNA structures showed unresolved electron densities around residues W15-17 (encompassing the flipped-out nucleotides) on both strands of the DNA (**Figures S3 & S4**). Despite the missing densities connecting them, however, the visible DNA segments clearly indicated that the DNA was kinked and unwound in a manner highly similar to that of the ‘open’ conformations (**Figures 3**). The observation of such similar ‘open-like’ conformations for both Δβ-hairpin2 and Δβ-hairpin3 mutants indicate that the majority of the ‘opening’ process does not require either β-hairpin2 or β-hairpin3 even though the ‘open-like’ DNA may entail greater conformational variabilities at the ‘open’ site.

**Figure 3.**
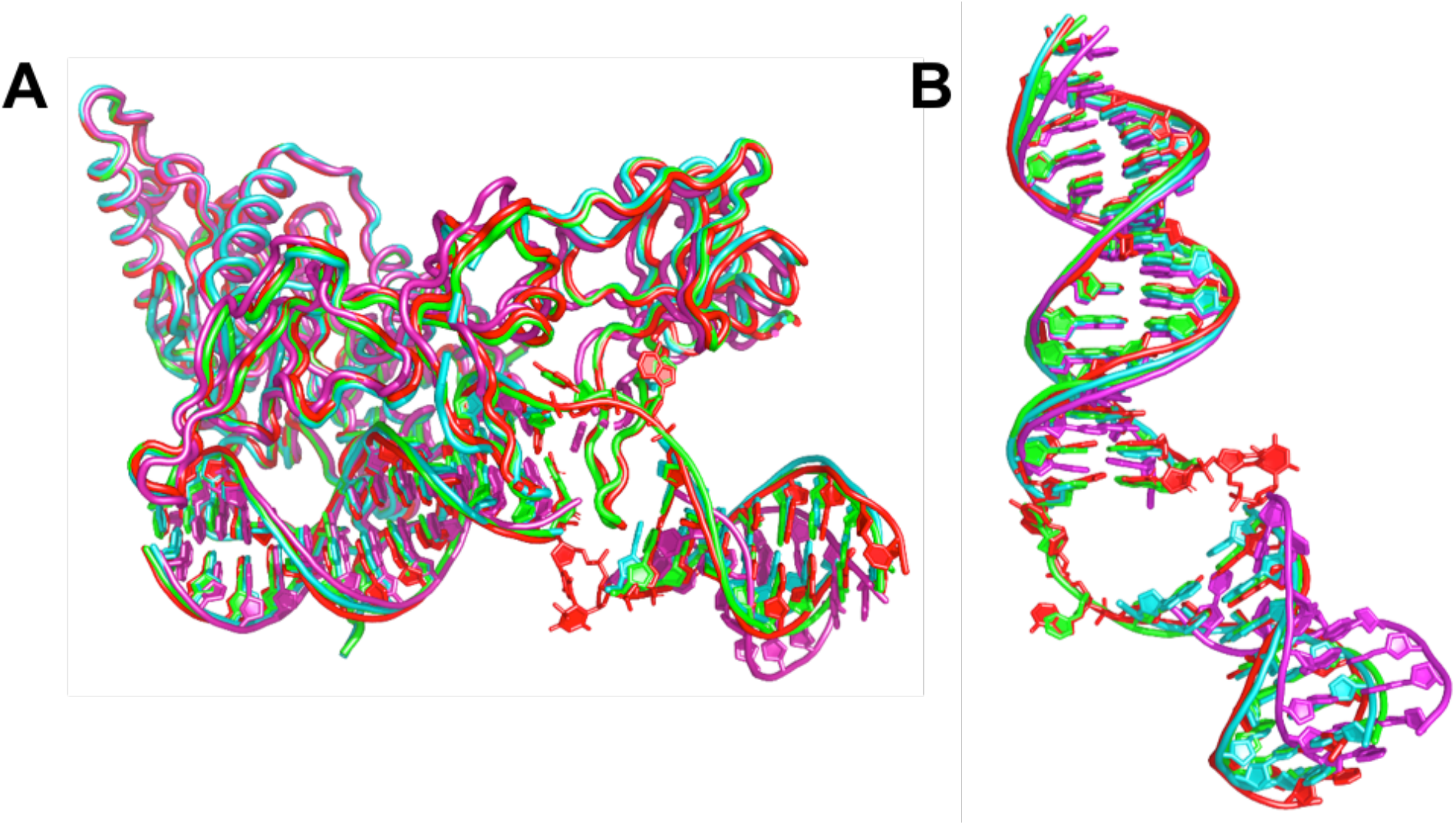
Comparison of the ‘open’ and ‘open-like’ crystal structures of Rad4–Rad23–DNA complexes. **(A)** Superposition of Rad4-Rad23-DNA crystal structures. The ‘open’ structures of WT Rad4 bound to 6-4PP (PDB ID: 6CFI) and of WT Rad4 tethered to CCC/GGG matched DNA (4YIR) are in red and green, respectively. The Δβ-hairpin3 and Δβ-hairpin2 mutants tethered to CCC/GGG matched DNA (cyan and magenta, respectively) show similar ‘open-like’ conformations. **(B)** Superposition of the DNA molecules extracted from the structures shown in (A). The difference in the DNA tails shown by the Δβ-hairpin2 (magenta) is partly due to the differences in crystal packing (**SI Discussion & Figure S2**).

### Rad4-tethered matched DNA shows a dynamic ‘open’ conformational landscape in solution

Although the crystallographic studies showed the 3D structures of the Rad4-DNA complexes in detail, such structures remain as snapshots captured in crystals, and whether the tethering-facilitated DNA ‘opening’ by Rad4 happens in solution remains to be examined. Also, the conformational states of molecules in solution can be much more complex and heterogeneous than depicted by crystal structures (43,44). Indeed, we previously showed such structural heterogeneity in CCC/CCC mismatched constructs without/with bound Rad4 and mapped out the population distributions of distinct DNA conformations that coexist in solution using fluorescence lifetime (FLT) analyses combined with a unique set of fluorescence resonance energy transfer (FRET) probes (tC^o^ and tC_nitro_) incorporated in DNA (34). The tC^o^ and tC_nitro_ are a FRET donor and acceptor pair, respectively (30,45). As cytosine analogs, these probes retain normal Watson–Crick pairing with guanines with minimal perturbation of DNA structure and stability (29,30) (**Figure 4A**). Furthermore, the rigid exocyclic ring and its base stacking interactions hold these nucleotide analogs in relatively fixed orientations within the DNA helical structure, making their FRET sensitive to subtle distortions in DNA helicity that alter the probes’ separation and/or relative orientation (31–33). For example, Rad4-induced untwisting and ‘opening’ of 3-bp mismatched DNA could be monitored by the FRET efficiency between tC^o^ (donor) and tC_nitro_ (acceptor) placed on either side of the mismatch (29,34). The FRET efficiency (E) relates directly to the lifetimes of the excited donor fluorophore, as 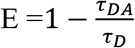, where *τ_DA_* and *τ_D_* are the donor lifetimes in the presence and absence of the acceptor, respectively. Also, by using the maximum entropy method (MEM) to analyze the fluorescence decay traces, one can robustly dissect multiple FRET components and their fractional amplitudes without having to *a priori* specify the number of lifetime exponents as in the more conventional discrete exponential analyses (38,39,46). The distinct FRET values and their fractional amplitudes obtained from each sample represent distinct co-existing conformations and their relative populations, respectively (34). The lifetime approach offers distinct advantages in obtaining FRET distributions over complementary approaches like single-molecule FRET, such as the ability to work with low quantum yield fluorescent probes and the superior (sub-nanoseconds) time-resolution that can capture distributions of rapidly fluctuating conformations (34,47).

**Figure 4.**
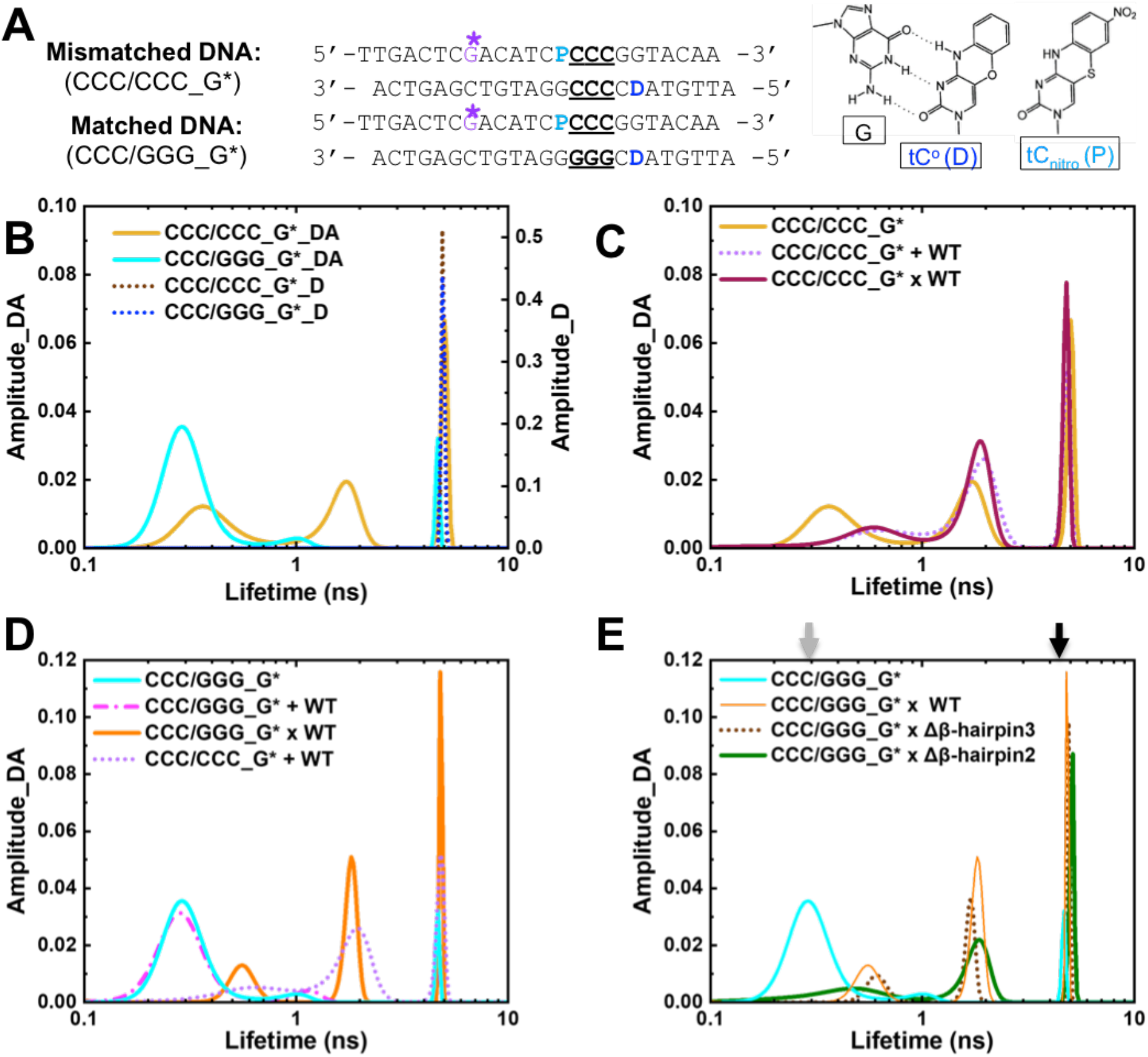
DNA conformational landscapes in solution obtained by fluorescence lifetime measurements. **(A)** DNA constructs for FLT studies. “D” indicates tC^o^ (FRET donor) and “P” is tC_nitro_ (FRET acceptor). G* is disulfide-modified guanine for tethering. Positions of the 3-bp CCC/CCC mismatches or CCC/GGG matched sequences are underlined. (*right*) Chemical structures of tC^o^ and tC_nitro_ and Watson-Crick type base pairing of tC^o^ with a guanine (G). **(B-E)** Representative FLT distributions from MEM analyses for different DNA and protein-DNA complexes. “_D” indicate DNA with donor only; “_DA” indicate DNA with donor/acceptor pair. Results in (C-E) are all from DNA_DA. **(B)** FLT distributions of mismatched CCC/CCC_G*_DA (yellow) or _D (dotted brown) and CCC/GGG_G*_DA (cyan) or _D (dotted blue) **(C)** FLT distributions of mismatched CCC/CCC_G* DNA when by itself (yellow), noncovalently bound to (“+”; dotted violet) or site-specifically tethered with (“x”; deep purple) WT Rad4. **(D)** FLT distributions of matched CCC/GGG_G* DNA when by itself (cyan), noncovalently bound to (“+”; dotted magenta) or tethered with (“x”; orange) WT Rad4. The FLT distribution of CCC/CCC_G* + WT Rad4 is also shown in dotted violet. (**E)** FLT distributions of CCC/GGG_G* when by itself (cyan) or tethered to WT Rad4 (orange), Δβ-hairpin3 (dotted brown) and Δβ-hairpin2 (green). All amplitudes indicate the normalized, fractional amplitudes. The arrows indicate the lifetimes corresponding to the computed FRET efficiencies for B-DNA conformation (grey) and for the DNA conformation in the Rad4-bound ‘open’ crystal structure (black). Reproducibility of FLT distributions for each sample is shown in **Figure S6.** Full reports of the lifetimes, fractional amplitudes, FRET efficiencies of each peak as well as the sample’s average FRET efficiencies are in **Table S2**.

Here, we applied this approach to the tethered Rad4-DNA complexes for the first time. The tC^o^-tC_nitro_ probes were introduced to the CCC/CCC mismatched DNA or the CCC/GGG matched DNA on either side of the putative ‘open’ site as before (29,34) and a crosslinkable G* was introduced as done for crystallization (**Figures 4A & S5**). The fluorescence lifetime distributions of various DNA and DNA-protein samples are shown in **Figures 4 & S6**. In the absence of the acceptor (thus no FRET), both the matched and mismatched donor-only DNA duplexes (DNA_D) showed a single lifetime peak corresponding to the intrinsic lifetime of the donor fluorescence (τ_D_ : 4.9 ns)(**Figure 4B**, dotted blue and brown); this donor lifetime is also insensitive to the absence and presence of Rad4 (34). On the other hand, the matched DNA containing both the donor and acceptor probes (CCC/GGG_G*_DA) showed a major lifetime peak (τ_DA_) at 0.32 ns corresponding to a FRET efficiency (E) of 0.94, with a fractional amplitude (thus fractional population) of 86% (**Figure 4B & S7A**, cyan; **Table S2**), with two minor peaks at 1.1 ns (E = 0.78) and 4.4 ns (E = 0.11) each occupying ~8% fractional amplitudes. The FRET value corresponding to the major peak (0.94) closely matches the FRET computed for an ideal B-DNA conformation with the given probe positions (0.93) (**Figure S7A**)(34,48). Furthermore, the FLT profile of CCC/GGG_G* matched well with that of the CCC/GGG DNA without G* (in which G* is replaced with normal G) (**Figure S7B**). Altogether, these results confirm that the matched CCC/GGG_G* DNA adopts a predominantly B-DNA conformation like CCC/GGG and other matched DNA sequences without G* examined previously (34).

In contrast to the matched DNA, the mismatched CCC/CCC_G*_DA showed a heterogeneous profile in its FLT distribution with three peaks: 0.40 ns (39% in fractional amplitude), 1.7 ns (33%) and 5.2 ns (28%), corresponding to high (~0.9), medium (~0.6) and low/zero FRET (~0.0), respectively (**Figures 4B & S7A** yellow; **Table S2**). The profile was again fully consistent with the FLT of the CCC/CCC_DA without G* (**Figure S7B**). As also shown before (34), the high fractional amplitude of the DNA in the low/zero FRET peak was not due to an excess of free donor strand, as this population persisted even when the DNA duplex was annealed with 50% excess of the acceptor strand (**Figure S7C**). The heterogeneous profile reflects the high intrinsic deformability of the CCC/CCC mismatches with multiple coexisting conformations (34).

Next, we examined the DNA in the presence of noncovalently interacting, equimolar WT Rad4. The profile of the matched CCC/GGG_G* did not change when Rad4 was added (**Figure 4D**, cyan vs magenta) and retained the majority B-DNA conformation. Yet, the mismatched CCC/CCC_G* showed a shift in its distribution towards higher lifetimes (lower FRET) indicating more distorted DNA conformations, away from the B-DNA (high FRET) (34,48) (**Figure 4C**, yellow vs violet; **Table S2**). However, instead of presenting one major low/zero FRET peak corresponding to the conformation found in the Rad4-bound ‘open’ crystal structure (computed E=0.04), the profile showed multiple broad peaks as observed before with the same DNA without G* (34). These results consistently indicate highly heterogeneous conformational states for the CCC/CCC mismatches in solution, even when it is specifically bound to and ‘opened’ by Rad4 and underscore the robustness of our approach in capturing the conformational landscapes of DNA in solution.

Following these, we next examined the lifetimes of DNA covalently tethered to Rad4. First, the Rad4-tethered mismatched CCC/CCC_G* showed a profile closely resembling that of the ‘open’ DNA noncovalently bound WT Rad4 (**Figure 4C**, deep purple vs. violet). These results provide an important control showing that the site-specific crosslink did not alter the free energy landscape in solution, which underlies the distribution of natural, noncovalently formed ‘open’ conformations. In contrast, the profile for Rad4-tethered matched DNA (CCC/GGG_G*) deviated significantly from its free or Rad4-bound forms and showed a remarkable resemblance to the mismatched DNA (CCC/CCC_G*) when bound/tethered to Rad4, although the widths of each peak were narrower than those with the mismatched DNA (**Figure 4D**, orange vs. violet; **Table S2**). By eliminating the possibility of heterogeneous binding register using site-specific tethering, the results unequivocally establish that: (1) the tethered/stalled protein could ‘open’ the matched CCC/GGG DNA at the probed site to form a complex similar to that formed with mismatched DNA; (2) the heterogeneous lifetime profile is not due to heterogeneous binding registers but reflects an inherent property of the ‘open’ complex in solution. The multiple FLT-FRET peaks reflect the inherently heterogeneous DNA conformations, particularly in terms of untwisting and bending around the ‘open’ site as sensed by the FRET probes. Each conformation (represented by each peak), however, may be less broadly sampled around an average conformation in matched DNA compared with mismatched DNA.

Finally, we examined the conformational distributions of the Rad4 β-hairpin mutants tethered to CCC/GGG_G*. Both Δβ-hairpin2 and Δβ-hairpin3 showed heterogeneous profiles in the FLT distributions closely mimicking that of the WT Rad4 although WT and Δβ-hairpin3 shared closer similarities to each other than to Δβ-hairpin2 (**Figure 4E** & **Table S2**). To further confirm that the altered FLT profiles of CCC/GGG were due to the tethering, we treated the tethered Δβ-hairpin3 x CCC/GGG_G* DNA complex with the reducing agent, dithiothreitol (DTT) that abolishes the disulfide tethering. After DTT treatment, the FLT profile reverted to a profile close to that of the free DNA showing predominantly B-DNA (lifetime ~0.3 ns) (**Figure S8**). Thus, the ‘open’ complex formed with matched DNA could be detected only under tethered conditions and could be reversed once Rad4 was allowed to freely diffuse away. These results are consistent with our previous studies that showed that crystallization of Rad4-bound undamaged DNA required tethering, in part because nonspecific ‘opening’ events that occur at random are inherently heterogeneous and not spatially synchronized and thus difficult to detect in an ensemble experiment. As demonstrated here, this tethering strategy also overcomes this experimental limitation under solution conditions. Altogether, these results are consistent with the similarities observed among the ‘open’ and ‘open-like’ crystal structures and show that tethering-facilitated matched DNA ‘opening’ occurs in solution with the mutants similarly as with WT.

### MD simulations show that the two mutants promote DNA distortions towards ‘opening’ in distinct manners

The crystallographic and FLT-FRET studies described above showed that Rad4 can form similarly ‘open’ conformations even without the full β-hairpins of BHD2 or BHD3. To probe the roles of the β-hairpins further, we performed 2-μs all atom MD simulations on the initial binding process between Rad4 and the CCC/GGG DNA sequence bound at a single register. Our previous MD studies showed that the initial binding states between WT Rad4 and different DNA adducts/lesions can meaningfully capture the differences between the lesions efficiently ‘opened’ by Rad4/XPC and repaired by NER versus nonproductive binding that is resistant to ‘opening’ and NER (25,41). In the 2-μs simulations with the WT and the Δβ-hairpin2 and Δβ-hairpin3 mutants bound to CCC/GGG, stable BHD2 conformations were achieved at ~1 µs. Hence the 1–2 µs trajectories were taken as the initial binding state (IBS) ensembles for each complex (**Figure S9, Movies S1 and S2**). Various structural parameters were analyzed for the IBS, and current results were compared with our previous study for the untethered WT Rad4–6-4PP complex, as a representative system where Rad4-induces DNA ‘opening’ for a specific substrate (25).

First, we monitored the BHD2-occupied AlphaSpace (AS) volumes and the DNA untwist angles at the putative ‘open’ site. The AS volume measures the extent of BHD2 binding in the DNA minor groove. The previous MD studies with 6-4PP and various bulky organic adducts have shown this parameter to correlate well with the lesion recognition propensity (25,41). The computed AS volumes were 165 Å^3^ and 286 Å^3^ for the WT and Δβ-hairpin3 binding to the CCC/GGG duplex, respectively, compared to 349 Å^3^ for the WT initial binding to 6-4PP (25,49) (**Figure 5A,B**). The successful engagement of BHD2 caused the CCC/GGG DNA to untwist by 9 ± 4º for the WT and 15 ± 4º for the Δβ-hairpin3 mutant, while it was 27 ± 4º for the WT bound to the 6-4PP lesion (**Figures 5B and S10,** see Methods) (25). The greater extent of untwisting in WT–6-4PP is in part due to the flipping of the nucleotides in the bottom strand, opposite the 6-4PP lesion; no nucleotide flipping was seen in the simulations with the CCC/GGG DNA bound to either WT or Δβ-hairpin3. Compared with these values at the IBS, the untwist angle shown in the final ‘open’ crystal structure of the WT-bound 6-4PP DNA was 89º (25).

**Figure 5.**
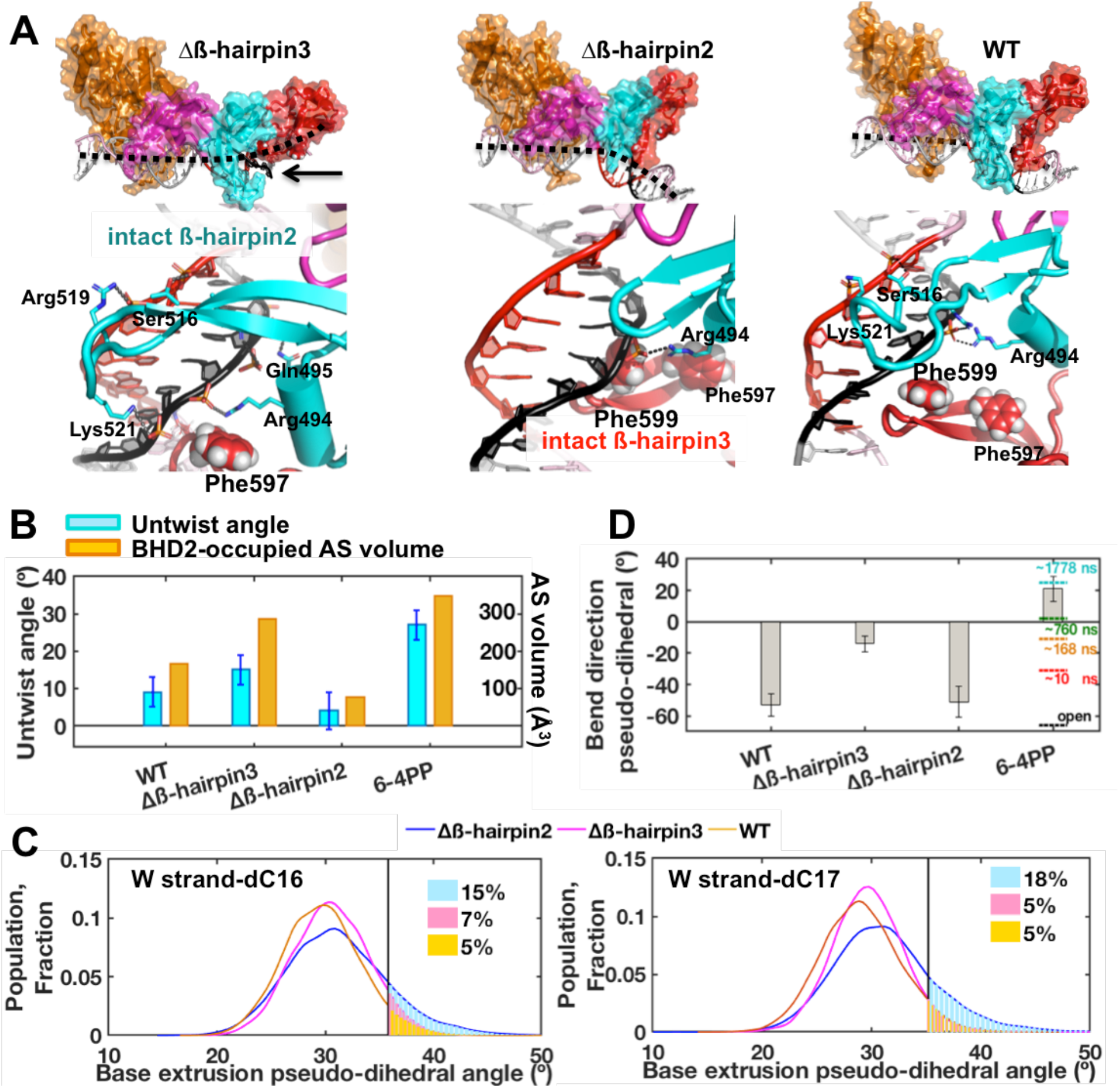
Initial binding states of the β-hairpin mutants with the CCC/GGG duplex obtained by MD simulations. **(A)** *(top)* Best representative structures of the initial binding states (IBS) show different effects of β-hairpin truncations. Black dashed lines show DNA bend directions (also see panel D); the minor groove around the GGG sequence (red) is below the dashed lines. *(bottom)* Enlarged views from the minor groove side. For Δβ-hairpin3, the intact β-hairpin2 inserts further into the minor groove than in the WT to promote untwisting. For Δβ-hairpin2, the intact β-hairpin3 approaches the potential flipping bases from the major groove more than in the WT. SI Movies S1 and S2 also display major groove views, which provide a good view of the BHD3 hairpin on that side. **(B)** Untwist angle and BHD2-occupied minor groove AS volume (48) show that untwisting correlates with extent of BHD2’s minor groove occupancy, both of which were more pronounced for Δβ-hairpin3 compared to Δβ-hairpin2. **(C)**Δβ-hairpin2 promotes extrusion of partner strand bases compared to WT and Δβ-hairpin3. The extrusions are facilitated by Phe599 in the β-hairpin3 that moved closer to DNA than in WT. **(D)** DNA bend directions show bending towards the minor groove (negative values) for CCC/GGG DNA, unlike with 6-4PP. The values of the bend directions for the 6-4PP along the MD trajectory (at different simulation times) are shown in color dashed lines and the value for the crystal structure of the open complex is in black dashed line (25). The standard deviations of block averaged means (80,81) for the untwist angles (B) and the bend direction pseudo-dihedral angles (D) are shown. Full details of the block averaging method are given in SI Methods.

For the Δβ-hairpin2-DNA complex, BHD2 formed one hydrogen bond to the DNA backbone around the putative ‘open’ site containing the CCC/GGG sequence and stayed stably bound (**Figure 5A**). However, due to the shortened β-hairpin2, BHD2 failed to engage with the minor groove (AS volume of 77 Å^3^); as a result, the DNA untwisting was very modest (4 ± 5º) compared to the ones observed with WT and Δβ-hairpin3 (**Figure 5B**). Instead, the intact β-hairpin3 exhibited enhanced dynamical interactions with the major groove, in which Phe599 moved closer to the partner strand bases than in the WT case because β-hairpin2 is truncated and causes much less untwisting (**Figures 5A and S10**). Phe599 is the first Phe along the partner base flipping pathway previously identified in Rad4 (50); it is a crucial residue for achieving productive opening of DNA duplexes, by directing partner strand base flipping and guiding β-hairpin3 insertion. The Phe599 interaction promoted the dynamic base extrusions near the β-hairpin3 (W16-17 cytosines, **Figure 2A**), as tracked by the base extrusion pseudo-dihedral angles along the trajectory (**Figures 5C and S11**). Such evidence of episodic partial extrusion was essentially absent in the WT and Δβ-hairpin3 mutant, let alone the unbound DNA (**Figure S11**). We found that this episodic partial extrusion is supported by van der Waals interactions between the flippable cytosine and Phe599, and hydrogen bonding of the partner guanine with Arg601 and Ser603 at the hairpin tip of BHD3 (**Figure S12**). Enhanced base extrusion thus provides a mechanism by which the Δβ-hairpin2 enables the ‘open’ conformation on the DNA despite lacking the β-hairpin2.

DNA in our various complexes exhibited similar bend angles (**Figure S10**) but differed in their bend directions, which may also change during the binding process (25). In order to monitor the DNA bending directions, we have computed a bend direction pseudo-dihedral angle (see **Figure S10** for definition), as a general indicator of bending toward major or minor groove: this parameter differs when bending is toward the major or toward the minor groove and is more negative when bending is toward the minor groove. Bending toward the minor groove can be promoted by protein–DNA interactions on the major groove side (*e.g.*, β-hairpin3). This was revealed in our analysis of the bend direction from our previous simulation for the WT-6-4PP initial binding (25). The bend direction showed progressive changes as the simulation advanced, starting from −31º; it became maximally positive (21 ± 8º) after an adenine base complementary to 6-4PP was flipped out into the BHD2/3 groove and the DNA bent more towards the major groove **(Figure 5D)**. The value from the final ‘open’ crystal structure was −66º. Therefore, the DNA first bends progressively towards the major groove during the initial binding but later towards the minor groove in the final ‘open’ conformation. In our present cases, there was no nucleotide flipping during the initial binding, and the bend direction pseudo-dihedral angles remained negative on average throughout the simulation: −14 ± 5º for Δβ-hairpin3, −51 ± 10º for Δβ-hairpin2 and −53 ± 7º for WT-CCC/GGG (27) (**Figures 5D and S10**). For Δβ-hairpin3, the DNA bent more toward the major groove (less negative), as the truncation of β-hairpin3 allowed more extensive binding of the intact β-hairpin2 with the minor groove; this was also indicated by the AS volume analysis described above. For Δβ-hairpin2 and WT, the DNA bent more toward the minor groove (more negative) as the remaining β-hairpin3 engaged DNA from the major groove side. The results unveil the distinct roles of the two β-hairpins in steering the bend directions during the DNA ‘opening’ process.

It is also interesting to note that the current simulation of WT with CCC/GGG DNA showed differences from the previous simulation of WT on the 6-4PP lesion. For instance, the degree of DNA untwist and BHD2-associated AS volume were smaller than with 6-4PP. Also, the bend direction analysis showed significant bending towards the minor groove, unlike that observed with 6-4PP. The results indicate that even for the same WT protein, the binding trajectories may differ depending on the DNA substrate. The concept that different DNA substrates can have distinct binding trajectories is consistent with our previous studies performed with DNA lesions whose repair propensities varied (25,41,50,51).

Altogether, the MD results show the key role of each hairpin when the other is truncated. Most strikingly, we show that β-hairpin2, by binding in the minor groove, facilitates untwisting and bending more toward the major groove, while β-hairpin3 facilitates partner base extrusion and bending more toward the minor groove. A combination of untwisting, bending, and base flipping in the DNA achieved through the interplay of the β-hairpins ultimately drives the formation of the ‘open’ structure.

## DISCUSSION

### Implication of tethering-facilitated DNA ‘opening’ in the DNA lesion recognition by Rad4/XPC

How XPC/Rad4 can efficiently scan and recognize a wide variety of DNA lesions and thus enable versatile NER has been a central question. Previous studies suggested that (1) lesion recognition by Rad4/XPC occurs by indirectly sensing local destabilization in a DNA duplex commonly resulting from DNA damage and selectively ‘opening’ those DNA sites; (2) the selective ‘opening’ is kinetically controlled, as determined by the balance between the free energy barrier to ‘open’ a DNA site (thus reflected in the protein-induced DNA ‘opening’ time) and the free energy barrier to diffuse away from that site. In this ‘kinetic gating’ model, Rad4/XPC does not rely on any specific structural feature of a lesion but can work to locate diverse lesions including 6-4PP and various bulky DNA adducts as it searches along DNA using thermal energy alone (*i.e.,* no chemical energy expenditure such as ATP hydrolysis). The experimentally measured Rad4-induced DNA ‘opening’ time and the residence time of Rad4 per DNA site are consistent with this model: laser temperature-jump perturbation spectroscopy (T-jump) experiments showed that the Rad4-induced mismatched DNA ‘opening’ takes several milliseconds while the protein’s nonspecific untwisting interrogation happens in 100-500 μs time scales (27,29). In comparison, the protein’s residence time during nonspecific searching by 1-D diffusion is ~1-600 μs per base pair, as measured by single-molecule microscopy (28) while the Rad4-induced ‘opening’ of undamaged DNA is expected to be much slower. Therefore, with such limited residence time, it is unlikely that Rad4/XPC may fully ‘open’ undamaged DNA although it may nonspecifically untwist DNA as it moves along. However, when the residence time is prolonged, for instance by tethering, even the undamaged DNA could be ‘opened’ as previously indicated by a crystal structure of WT tethered to undamaged CCC/GGG DNA (27). Nonetheless, the evidence for such tethering-facilitated DNA ‘opening’ of undamaged DNA was limited to the crystal structure and had not been reported for complexes in solution, although previous AFM studies showed that the degrees of DNA bending were similar between Rad4-bound damaged DNA versus Rad4-bound undamaged DNA (27).

Here, we applied the tC^o^-tC_nitro_-based FLT-FRET approach to examine the solution conformations accessible to Rad4-DNA complexes when tethered to undamaged DNA. The broad range of conformations observed for the ‘open(-like)’ complexes indicate varying degrees of untwisting and bending in the DNA coexisting in solution. Similar profiles were observed for both the Rad4-tethered matched DNA and the untethered yet specifically recognized mismatched DNA, despite the fact that unbound matched DNA exhibited on average a homogeneous B-DNA while the unbound mismatched DNA could adopt varied heterogeneous conformations by itself; these results indicate that the dynamically fluctuating and flexible DNA conformations in the Rad4-induced ‘open’ complexes are an inherent feature of the ‘opened’ DNA rather than originating from the intrinsic dynamics of the DNA alone. We posit that this flexible characteristic of Rad4-bound ‘open’ DNA may be a key feature for the downstream NER in which the strand-separated DNA bubble needs to be further expanded up to ~10-20 bp for lesion verification through a multistep process involving various NER factors (e.g., XPD, XPB and XPA) (52,53): flexibly ‘open’ DNA when bound to Rad4, as observed in this and our previous study (34), may be required as a DNA platform amenable to further downstream processing of NER.

In a broader sense, the DNA ‘opening’ of a given site facilitated by prolonged residence time at that site may help explain the mechanisms of XPC within and beyond NER that cannot be explained solely by lesion binding preferences of XPC (57–64). In these cases, proteins interacting with XPC (e.g., UV-DDB or DNA glycosylases or PARP or transcription factors) and/or post-translational modifications on XPC could help ‘stall’ the protein on DNA and induce opening of otherwise non-cognate DNA. Such ‘kinetic gating’ also has broad implications as it may generally apply to various other site-specific DNA-binding proteins as a way to help reduce wasteful interrogation at each and every site.

### Role of β-hairpin2 and β-hairpin3 in the DNA lesion recognition by Rad4/XPC

While it has been shown that DNA ‘opening’ by Rad4/XPC involves four consecutive domains including three structurally related β-hairpin-containing BHD domains, the roles of the β-hairpins in ‘opening’ DNA has been unclear. Here, we examined the role of β-hairpins in BHD2 or BHD3 in ‘opening’ undamaged DNA using a combination of x-ray crystallography and fluorescence lifetime-based conformational analyses on tethered complexes, with further insights gained from MD simulations. Our study, as depicted in a schematic summary in **Figure 6**, shows that (1) mutants lacking either β-hairpin can undergo the bulk of the ‘opening’ process on matched CCC/GGG DNA under tethered conditions similar to the WT protein under the same condition; (2) β-hairpins2/3, however, are still important for stably holding the two flipped-out nucleotides in the BHD2/3 groove; (3) the conformational landscapes of the ‘open’ complex with WT or the ‘open-like’ complex with the mutants are intrinsically dynamic around the ‘open’ DNA site and are similar to each other in solution, consistent with the crystal structures; (4) despite the similarities in the DNA conformations adopted with the two mutants and WT, atomistic MD simulations indicate that the β–hairpins may promote DNA ‘opening’ in distinct manners and may compensate for each other when one is lacking.

**Figure 6.**
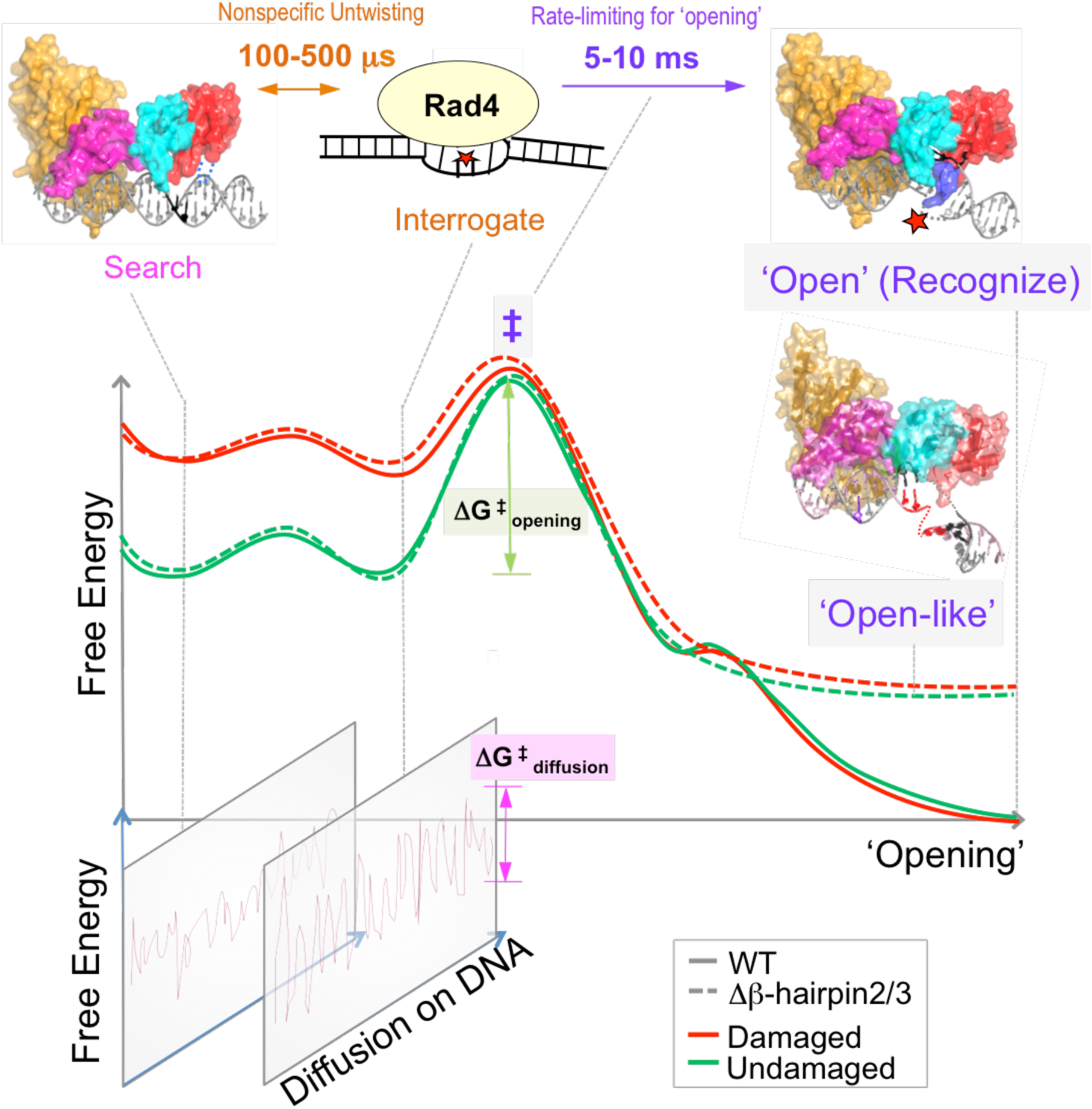
Proposed DNA ‘opening’ trajectory and ‘kinetic gating’ mechanism of Rad4/XPC. The top panel illustrates distinct binding modes for Rad4/XPC as it searches for, interrogates, and recognizes a damaged site, and the time scales for fluctuations between these modes, based on prior studies (18–20). The middle panel shows a schematic free-energy profile along the ‘opening’ trajectory. The faster 100- to 500-µs nonspecific untwisting step entails a smaller energetic barrier than the slower 5- to 10-ms rate-limiting step (‡) of the ‘opening’ process. The rate-limiting step involves sufficiently unwound and bent DNA but with the nucleotides not yet fully or stably flipped out into the BHD2/BHD3 groove (19). The free energy barrier (ΔG^‡^_opening_) for ‘opening’ damaged DNA (red) is naturally lower than that for undamaged DNA (green) as DNA damage destabilizes the B-DNA structure. For Rad4 mutants that are lacking either β-hairpin2 or β-hairpin3, the protein can still overcome ΔG^‡^_opening_ to form ‘open-like’ structures that exhibit the same extent of unwinding as the fully ‘open’ structure with the WT (18), as monitored by the tC^o^-tC_nitro_ FRET probes; however, the ‘open-like’ structures with the mutants are less stable and do not show well-resolved flipped-out nucleotides in the crystal structures (this study). MD simulations indicate that there can be more than one pathway that leads to ‘open-like’ or ‘open’ structures and demonstrate that the two β-hairpins function in a concerted manner to promote ‘opening’, but can also compensate for each other when one β-hairpin is lacking. The bottom panel illustrates that for each step along the ‘opening’ trajectory, there is also a kinetically competing process of diffusion of Rad4/XPC along the DNA, characterized by ΔG^‡^_diffusion_. For undamaged DNA, the high ΔG^‡^_opening_ compared with ΔG^‡^_diffusion_ favors the protein diffusing away before ‘opening’ a given site, while for damaged DNA this competition favors ‘opening’. However, when tethered, the diffusion of the protein is blocked and it can ‘open’ that site as long as the ΔG^‡^_opening_ is thermally surmountable. Our study suggests that stalling of Rad4/XPC by another protein may be a mechanism employed by NER to enable the ‘opening’ of more resistant NER lesions.

Our results also add to the similarities and differences between WT and Δβ-hairpin3 previously noted under non-tethered conditions. Previous T-jump studies concluded that Δβ-hairpin3 or ΔBHD3 (lacking the entire BHD3 domain) could exert fast nonspecific untwisting on DNA (100-500 μs) as a way to interrogate DNA. However, the slow kinetics (5-10 ms) corresponding to the rate-limiting step of the ‘opening’ could not be directly measured for Δβ-hairpin3 and was observed for ΔBHD3 only at higher temperatures (29). The diffusional movements of Rad4 on UV-damaged λ-DNA tightropes were also examined using single molecule imaging (28): while Δβ-hairpin3 had a larger fractional population of proteins that was motile and diffused freely along the DNA rather than being immobilized (presumably at a lesion site) compared with WT, both the mutant and WT could similarly localize to and stably associate with damaged DNA containing fluorescein-dT lesions. Notably, the UV survival and CPD repair rates in yeast found little difference between the WT and the Δβ-hairpin3 strains (28). Thus, despite the diminished ability of Δβ-hairpin3 to localize to and fully ‘open’ DNA lesions *in vitro*, the β-hairpin3 may be generally dispensable for the Rad4/XPC function in cells. It is interesting to note that the human XPC mutant lacking the entire β-hairpin3 (residues 789-815 corresponding to 589-616 in Rad4) failed to localize to local UV-irradiated areas in cells on its own but required UV-DDB for the localization (54). It is tempting to speculate that UV-DDB may play a role analogous to tethering in our study and enable DNA ‘opening’ by the XPC mutant by helping its recruitment to and preventing its diffusion from UV lesions. The impact of Δβ-hairpin3 in yeast may also have to account for other interacting factors such as Rad7-Rad16 that may function equivalently to UV-DDB in human cells (55,56). Finally, our results also suggest that the impact of Δβ-hairpin2 may be similar to that of Δβ-hairpin3 in cellular or biophysical conditions.

Several other NER proteins also feature a β-hairpin that functions in DNA-binding (65). For instance, UvrB interacts with DNA by inserting its β-hairpin between the strands of the duplex DNA, locking one of the strands between the β-hairpin and another domain (66). Deletion of the β-hairpin resulted in a decrease in the lifetime and increase in the 1-D diffusion rate on DNA for the UvrBC complex (67). XPA or Rad14 in yeast also possess a β-hairpin involved in DNA binding although, in this case, the β-hairpin is involved in packing against the exposed end of a DNA duplex with or without a lesion using a conserved aromatic residue (Trp175 in XPA; Phe262 in Rad14) (68–72). Mutation of Phe262 to alanine abolished the DNA binding of Rad14 (69). While these β-hairpins in UvrB and Rad14/XPA play a role in DNA binding in NER, β-hairpins 2/3 in Rad4 seem distinct in that they appear in tandem within one protein and that they can complement each other. It is noteworthy that most *XPC* patient mutations result in various truncation mutations that result in the absence of XPC protein detectible from cell extracts by Western blots, RT-PCR or activity assays (73–80). The complementary roles of the β-hairpins revealed in this study may extend to other domains such as BHD1, and we speculate that the relative paucity of point mutations in *XPC* patients may be at least in part due to high cooperativity and complementarity among different protein domains/motifs. Many of these important unresolved questions merit further investigation.

In conclusion, our work represents the first crystallographic and MD simulation studies of any Rad4/XPC mutant bound to DNA, revealing novel insights into the roles of the β-hairpins in lesion recognition by NER proteins. The fluorescence lifetime studies in combination with the tethering strategy also provide the first direct evidence in solution for ‘opening’ of matched DNA by Rad4/XPC in support of ‘kinetic gating' and reaffirm the dynamic nature of the Rad4-DNA ‘open’ complexes in solution. Finally, our work showcases how the FLT-FRET conformational analyses can reveal complex structural landscapes in solution, and thus be highly complementary to ensemble-averaged structural studies with techniques such as x-ray crystallography and cryo-electron microscopy as well as single molecule FRET studies.

## Acknowledgements

We thank the staff scientists of LS-CAT in APS for their help in data collection. We also thank the members of the Min, Broyde and Ansari groups.

## Accession Number

Atomic coordinates and structure factors for the reported crystal structures of the Δβ-hairpin3-CCC/GGG and Δβ-hairpin2-CCC/GGG complex and have been deposited with the Protein Data bank under accession number 6UBF and 6UIN respectively.

