## Supporting Information for "Tethering-facilitated DNA ‘opening’ and complementary roles of β-hairpin motifs in the Rad4/XPC DNA damage sensor protein"

Contains:

SI Methods; SI Results; SI Discussion

SI Tables S1-S2

SI Figures S1-S12

SI Movies S1-S2

**SI Methods**

**Experimental setup for fluorescence lifetime measurements**. The fluorescence decays of DNA labeled with $tC^o$ and $tC_{nitro}$ were measured using a time-correlated single-photon counting (TCSPC) system (DeltaFlex, HORIBA) equipped with a Ti-sapphire laser source with a tunable range from 690 to 1040 nm (Mai Tai HP, Spectra-Physics). The fundamental beam (730 nm) was passed through a half-wave plate and then to a pulse picker for reducing the repetition rate of the beam from 80 MHz to 4 MHz. For the second harmonic generation, the fundamental 730 nm beam coming from the pulse picker was focused onto a thin β-barium borate (BBO) crystal in second-harmonic generator (Minioptics, Inc., Arcadia, CA), which generated a frequency-doubled beam centered at 365 nm. The remaining fundamental beam and frequency-doubled 365 nm beam were separated through a dichroic mirror. The fundamental pulse was given a time delay through an optical delay cable connected with a delay box. The residual fundamental was allowed to pass through a photodiode for triggering the start signal in DeltaFlex collection system. For excitation of $tC^o$, the laser pulses were passed through a monochromator set at 365 nm (band pass 10 nm) attached with the DeltaFlex set up. We used neutral density filter (FSQ-ND20, broadband UV-grade fused silica metallic filter, Newport corporation) to reduce power of the excitation light suitable for TCSPC measurements. The fluorescence from the samples passed through a long pass filter cut at 375 nm and the signal was collected at magic angle with the emission polarizer oriented 54.7 ° from the direction of the excitation polarizer, chosen as vertical. This magic angle configuration removes any depolarization effects in the measured fluorescence decay curves from the rotational dynamics of the labeled biomolecules that could be present when exciting with polarized laser pulses, and is the configuration recommended by Horiba for their fluorescence lifetime setups. The resulting fluorescence signal was entered through the entrance slit of emission monochromator set at 470 nm (band pass 10 nm) and collected by a Picosecond Photon Detection module (PPD-850, Horiba). The instrument response function (IRF) of the system was measured using a dilute aqueous solution (3% w/w) of LUDOX AM colloidal silica (Sigma-Aldrich). The full-width half maxima (FWHM) was ~425 ps. Fluorescence decay curves were recorded on a 100 ns timescale, resolved into 4,096 channels, to a total of 10,000 counts in the peak channel. The power of the laser delivered to the samples was 0.21 $mW/cm^2$ as measured by a General Tools Digital UVA/UVB Meter, 280-400 nm (#UV513AB).

**Initial models, water box and counterions for MD simulations.** The two Rad4 Δβ−hairpin mutants were prepared by removing either residues 515-527 in β−hairpin2 or residues 599-605 in β−hairpin3 from the previous WT Rad4 structure, which was based on the crystal structure of unbound Rad4, and connecting the end residues of the truncated β−strands with a peptide bond. The complexes of the Rad4 mutants docked to CCC/GGG DNA were prepared as in the 'docking model' of our earlier work (1), where the BHD2 and BHD3 domains are close to but not yet bound to the duplex. All molecular modeling was carried out using Discovery Studio 2.5 (Accelrys Software Inc.). The complexes were neutralized with $Na^+$ counterions and solvated with explicit TIP3P water (2) in a cubic periodic box with side length of 125.0 Å using the tLEAP module of the AMBER16 suite of programs (3).

**MD simulations.** We used the ff14SB force field for the MD simulations (4). All MD simulations were carried out using the AMBER16 suite of programs (3). The Particle-Mesh Ewald method (5) with 9.0 Å cutoff for the non-bonded interactions was used in the energy minimizations and MD simulations. Minimizations were carried out in three stages. First, 500 steps of steepest descent minimization followed by 500 cycles of conjugate gradient minimization were conducted for the water molecules and counterions with a restraint force constant of 50 kcal/(mol·$Å^2$) on the solute molecules. Then, 500 steps of steepest descent minimization followed by 500 cycles of conjugate gradient minimization were carried out for the water molecules and counterions with a restraint force constant of 10 kcal/(mol·$Å^2$) on the solute molecules. In the last round, 500 steps of steepest descent minimization followed by 500 cycles of conjugate gradient minimization were carried out on the whole system without restraints. The minimized structures were then subjected to three rounds of equilibration. First, each system was equilibrated at constant temperature of 10 K for 30 ps with the solute molecules fixed with a restraint force constant of 50 kcal/(mol·$Å^2$). Then the system was heated from 10 K to 300 K over 300 ps with the solute molecules fixed with a restraint force constant of 50 kcal/(mol·$Å^2$) at constant volume. In the last round of equilibration, the restraint force constant on the solute was reduced through three steps: at 10 kcal/(mol·$Å^2$) for 200 ps, at 1 kcal/(mol·$Å^2$) for 200 ps, and then at 0.1 kcal/(mol·$Å^2$) for 200 ps with constant pressure at 300 K. Following equilibration, production MD simulations for each system were carried out in a constant-temperature, constant-pressure (NPT) ensemble at 300 K and constant pressure of 1 Atm for 2 μs. The temperature was controlled with a Langevin thermostat (6) with a 5 $ps^{-1}$ collision frequency. The pressure was maintained with the Berendsen coupling method (7). A 2.0 fs time step and the SHAKE algorithm (8) were applied in all MD simulations. A 1 kcal/mol restraint was applied to the end base pair hydrogen bond donor and acceptor atom pairs during the production MDs. During the production MD of the Δβ–hairpin3-DNA complex, and an extra 5 kcal/mol constraint was also applied between Lys350 backbone C and dG3 (W strand) backbone P atoms (**Figure 2** in the main text) to 'tether' the TGD domain to the DNA duplex.

**Structural analyses**

***Principal component analysis (PCA) and RMSD analysis.*** The structures along each trajectory were clustered using the principal component analysis (PCA) method in the Bio3D package (9). PCA calculations were performed for the heavy atoms of the potential 'open' site 6-mer (nucleotide steps W14 – 19, **Figure 2A**) and the protein backbone atoms of BHD2. The clustering was performed with 2001 frames selected every 1 ns from each 2 μs trajectory. The structures from each ensemble were superposed to the first frame at the heavy atoms of the base pairs that are non-specifically bound to TGD and BHD1 (nucleotide steps W4–12, **Figure 2A**) and serve as our reference for structural changes around the lesion site. The principal components (PCs) represent different proportions of variance for the structural ensemble and are ranked from highest to lowest. Hierarchical clustering of structures in PC space was performed along the first seven PCs, which represent ~ 85% of the structural dynamics in the MD trajectories, using R (10). The clustering results are given in **Figure S9**. Each cluster is an ensemble of structures that exhibits similar structural dynamics and represents a sub-stage upon Rad4 initial binding to the duplex. The RMSD for the heavy atoms of the BHD2 domain and potential 'open' site 6-mer DNA were calculated for the ensemble of structures in the production MD. The best representative structure for each ensemble is defined as the one frame that has the shortest RMSD for the heavy atoms used in the calculation of RMSD values to all other frames.

***The AlphaSpace volume analyses for BHD2 binding to the minor groove.*** The best representative structure for the initial binding state of each complex was further analyzed to quantify BHD2's binding into the DNA minor groove around the potential 'open' site. The AlphaSpace (AS) volumes of the binding pockets in the DNA and their occupancies by BHD2 were calculated using AlphaSpace v1.0 (11): the DNA duplex was set as the receptor and BHD2 was set as the ligand. The total occupied AS volume was used to quantify the extent of BHD2 binding into the DNA minor groove. The value reflects the curvature and surface area of the DNA minor groove region that is occupied by BHD2.

***Analysis of DNA untwisting upon the binding of the Rad4 mutants.*** The untwisting of the DNA duplex along the initial binding trajectory was monitored by the Untwist angle, defined as Untwist = Twist $_{initial}$ –Twist. The twist angle was measured between the end base pairs of the potential 'open' site 6-mer (W14 and W19, **Figure 2A**) using the cpptraj module of AMBER16 (3). Twist $_{initial}$ is the ensemble average twist angle of the 6-mer during the first 1 ns of production MD, during which significant untwisting was not observed (**Figure S10**); this ensemble represents the state of the duplex before the engagement of Rad4, especially BHD2. Positive values indicate further untwisting and negative values indicate further twisting.

***DNA bend angles and bend direction pseudo-dihedral angles.*** We measured the helix bend angle for the duplex for nucleotide steps W11–21 (**Figure 2A**) using Curves+ (12) (**Figure S10**). The DNA's bend direction was measured using a pseudo-dihedral angle (defined in **Figure S10**) using the cpptraj module of AMBER16 (3).

***Distance between Phe599 and partner strand bases at the potential 'open' site.*** Phe599 is the first Phe along the partner base flipping pathway identified previously in Rad4 (13), and is a crucial residue for achieving productive opening of DNA duplexes, by directing partner strand base flipping and guiding β–hairpin3 insertion. The distance between Phe599 and the partner strand bases at the potential 'open' site was measured between the COM of the phenyl ring heavy atoms in Phe599 and the COM of the two cytosines (W16 and W17, **Figure 2A**) that would flip to bind to BHD2/3 in the 'open' structure, using the cpptraj module of AMBER16 (3) (**Figure S10D**).

***Base extrusion pseudo-dihedral angles.*** The base extrusions of partner strand nucleotides at the potential open site (dC's in W16-W17, **Figure 2A**) were monitored using a pseudo-dihedral angle, defined by four centers of masses (COMs): the COM of the C base heavy atoms, the COM of dC sugar ring heavy atoms, the COM of its 3' nucleotide's sugar ring heavy atoms, and the COM of its 3' base pair heavy atoms (**Figure S11**). The pseudo-dihedral angle was measured using the cpptraj module of AMBER16 (3). This dihedral angle is more positive when extrusion is via the major groove.

***Van der Waals interaction energy between BHD2/3 and partner strand nucleotides.*** The van der Waals interaction energy between the BHD2/3 amino acid side chains and each of the deoxycytidines at the potential 'open' site (W16 and W17, **Figure 2A**) was calculated using the cpptraj module of AMBER16 (3) for the Lennard-Jones potential **(Figure S12)**.

***Analysis of hydrogen bonding between BHD2/3 and DNA.*** Hydrogen bonds between BHD2/3 and the DNA duplexes were counted using each pair of donor and acceptor atoms, when they have a hydrogen bond (heavy-light-heavy atom) angle ≥ 145° and heavy-to-heavy atom distance ≤ 3.3 Å **(Figure S12)**.

***Block average analyses.*** The DNA structural parameters for the stable ensemble of each MD simulation were analyzed using the block averaging method (14,15). In brief, the time series data were divided into "blocks" with a block size that exceeds the longest correlation time, 50 ns in our case. The average for each block was computed and termed 'block average'. The mean values and the standard deviations of the block averages given in the main text were used to represent the average and the variance of averages.

Molecular structures were rendered using PyMOL 1.3.x (Schrodinger, LLC.). All MD simulation data were plotted using MATLAB 7.10.0 (The MathWorks, Inc.)

### SI Results

In further support of the results for our 2 μs simulations, we have now performed 30 additional 100 ns simulations, 10 for each of the three systems, started from the best representative structures (**Figure 5A** in the main text and **Figure S9**), with re-equilibration. Our results showed that the untwist angle remained within the standard deviation of the block averaged values from the long run. The values from the long runs (**Figure 5B** in the main text) are 9 ± 4º for the WT, 15 ± 4º for the Δβ-hairpin3 mutant, and 4 ± 5º for the Δβ-hairpin2 mutant. For the multiple runs, they are 10 ± 3º, 11 ± 3º, and 5 ± 4º. Hence the additional sampling with multiple runs supports the results obtained with the long runs.

### SI Discussion

**Crystal packing and structural discrepancies between the Δβ−hairpin2 and Δβ−hairpin3-DNA complexes.** While the 'open-like' structures formed by the two mutants closely resembled each other and were similar to the 'open' structure by the WT, some differences could be noted. The crystal containing Δβ-hairpin3 tethered to the CCC/GGG DNA belonged to the tetragonal space group $P4_12_12$ with one monomer in the asymmetric unit and diffracted to 4.7 Å with a Matthews coefficient of 3.83, similar to the previous crystal with the WT Rad4 (PDB ID: 4YIR). The crystal containing Δβ-hairpin2 tethered to the same DNA belonged to the hexagonal space group $P6_5$ with one monomer in the asymmetric unit and diffracted to 3.3 Å with a Matthews coefficient of 2.82. The tetragonal Δβ−hairpin3-DNA crystal had a z-axis that was almost twice as long as that of the hexagonal Δβ−hairpin2-DNA crystal (403 Å versus 264 Å) with slightly higher solvent content (68% versus 56%), indicating a less dense crystal packing. In both the crystals, the ends of the DNA were packed against another DNA molecule from the symmetry related molecules in a head-to-tail fashion (**Figure S2**). However, despite the higher resolution of the Δβ−hairpin2-DNA structure than the Δβ−hairpin3-DNA, Δβ−hairpin2-DNA had more nucleotides with missing electron densities than Δβ−hairpin3-DNA (four versus three). Furthermore, the DNA portion extended beyond BHD2/3 showed some difference between the Δβ−hairpin2 and the WT- and Δβ−hairpin3 crystals, likely due to the different crystal packing arrangement with neighboring molecules (**Figures 3B & S2**). For instance, the untwist angle defined as Untwist = Twist $_{initial}$ –Twist $_{crystal}$ (SI Method) was 94° for the DNA in the Δβ−hairpin3-DNA, very close to the untwist (89°) shown in the 'open' crystal structure of 6-4PP DNA bound to WT (16), whereas it was 72° for the Δβ−hairpin2-DNA structure. The distances between the backbone phosphorus atoms of the common visible nucleotides were 21.5 Å for top strand (W14 and W18) and 17.4 Å for the bottom strand (Y7 and Y12) in the Δβ−hairpin3-DNA complex; the equivalent distances were 24.4 Å and 16.4 Å in the Δβ−hairpin2-DNA complex (**Figure S4**). Altogether, these results showed that the Δβ−hairpin2-DNA showed slightly less unwound but more broadly disordered putative 'open' site compared with Δβ−hairpin3-DNA.

**SI Tables S1-S2**

**Table S1.** Data collection and refinement statistics (molecular replacement)

| PDB ID | **6UBF** (Δβ−hairpin3 x CCC/GGG) | **6UIN** (Δβ−hairpin2 x CCC/GGG) |
|---|---|---|
| **Data collection** | | |
| Wavelength (Å) | 0.9794 | 0.9794 |
| Space group | $P\ 4_12_12$ | $P\ 6_5$ |
| Cell dimensions | | |
| *a, b, c* (Å) | 78.7 78.7 403.8 | 77.6 77.6 264.0 |
| $\alpha, \beta, \gamma$ (°) | 90 90 90 | 90 90 90 |
| Resolution (Å) | 50.0 – 4.6 | 50.0 – 3.3 |
| $R_{sym}$ or $R_{merge}$ | 12.6 (39.7) | 9.1 (83.5) |
| $I / \sigma I$ | 23.2 (2.2) | 20.6 (2.7) |
| Completeness (%) | 100 (99.1) | 99.9 (99.4) |
| Redundancy | 13.7 (11.1) | 6.4 (6.4) |
| **Refinement** | | |
| Resolution (Å) | 48.7-4.6 (4.76-4.60) | 38.83-3.35 (3.47-3.35) |
| No. reflections | 17956 (1519) | 27956 (2519) |
| $R_{work}$ / $R_{free}$(%) | 32.3/33.8 | 20.7/25.8 |
| No. atoms | 5209 | 5197 |
| Protein | 4281 | 4280 |
| DNA | 928 | 917 |
| Water | 0 | 0 |
| *B*-factors (Å$^2$) | 331.29 | 139.57 |
| Protein | 62.3 | 56.3 |
| DNA (G47) | 91.3(91.7) | 88.3(91.7) |
| Water | 0 | 0 |
| R.m.s. deviations | | |

R.m.s., root mean squared.

*Values in parentheses are for highest-resolution shell.

**Table S2.** MEM fluorescence lifetimes and FRET efficiencies.

| | $\tau_1$ | $A_1$ (%) | $E_1$ | $\tau_2$ | $A_2$ (%) | $E_2$ | $\tau_3$ | $A_3{}^a$ (%) | $E_3{}^a$ | <E> |
|---|---|---|---|---|---|---|---|---|---|---|
| **CCC/CCC_G*** | 0.396 ± 0.018 | 39.28 ± 1.753 | 0.919 ± 0.003 | 1.723 ± 0.035 | 32.75 ± 4.187 | 0.649 ± 0.007 | 5.188 ± 0.237 | 27.97 ± 2.432 | -0.053 ± 0.049 | 0.560 ± 0.037 |
| **CCC/GGG_G*** | 0.316 ± 0.018 | 85.67 ± 1.331 | 0.935 ± 0.003 | 1.067 ± 0.177 | 8.040 ± 0.297 | 0.783 ± 0.036 | 4.374 ± 0.301 | 8.220 ± 1.038 | 0.110 ± 0.061 | 0.864 ± 0.000 |
| **CCC/CCC_G* + WT** | 0.975 ± 0.019 | 35.13 ± 1.088 | 0.802 ± 0.003 | 1.934 ± 0.002 | 41.81 ± 1.063 | 0.607 ± 0.000 | 4.807 ± 0.019 | 23.22 ± 0.258 | 0.024 ± 0.003 | 0.539 ± 0.000 |
| **CCC/GGG_G* + WT** | 0.327 ± 0.026 | 89.96 ± 6.033 | 0.933 ± 0.005 | 1.226 ± 0.181 | 5.477 ± 2.030 | 0.750 ± 0.036 | 4.653 ± 0.023 | 4.566 ± 3.996 | 0.054 ± 0.004 | 0.862 ± 0.000 |
| **CCC/CCC_G* X WT** | 0.728 ± 0.062 | 29.19 ± 1.968 | 0.852 ± 0.012 | 1.869 ± 0.005 | 43.69 ± 1.770 | 0.619 ± 0.001 | 4.801 ± 0.040 | 27.12 ± 0.199 | 0.024 ± 0.008 | 0.538 ± 0.000 |
| **CCC/GGG_G* X WT** | 0.608 ± 0.070 | 25.07 ± 1.210 | 0.876 ± 0.014 | 1.836 ± 0.009 | 46.85 ± 1.325 | 0.628 ± 0.001 | 4.827 ± 0.135 | 28.08 ± 0.657 | 0.188 ± 0.027 | 0.635 ± 0.006 |
| **CCC/GGG_G* X Δ$\beta$-hairpin2** | 0.567 ± 0.188 | 33.02 ± 1.707 | 0.884 ± 0.038 | 1.931 ± 0.074 | 34.54 ± 5.720 | 0.607 ± 0.015 | 5.197 ± 0.179 | 31.85 ± 4.936 | -0.055 ± 0.036 | 0.569 ± 0.061 |
| **CCC/GGG_G* X Δ$\beta$-hairpin 3** | 0.615 ± 0.011 | 24.79 ± 1.881 | 0.874 ± 0.002 | 1.676 ± 0.018 | 47.04 ± 4.402 | 0.659 ± 0.003 | 4.805 ± 0.181 | 28.17 ± 2.522 | 0.024 ± 0.036 | 0.697 ± 0.116 |

[a] Corresponding to the longest lifetime 'zero-FRET' component.

$\tau_n$, $A_n$, and $E_n$ (n=1,2,3) indicate the fluorescence lifetimes, normalized amplitudes and FRET efficiencies for each Gaussian peak of the sample containing both the donor and acceptor probes. The average FRET efficiencies of the samples were calculated as $< E >= 1 - \frac{<\tau_{DA}>}{\tau_D}$, where the donor-only lifetimes ($\tau_D$) for the DNA samples were 4.926 and 4.920 ns for CCC/CCC_G*_D and CCC/GGG_G*_D, respectively. The uncertainties reported are standard deviations (s.d.) from 2-3 independent sets of measurements.

**SI Figures S1-S12**

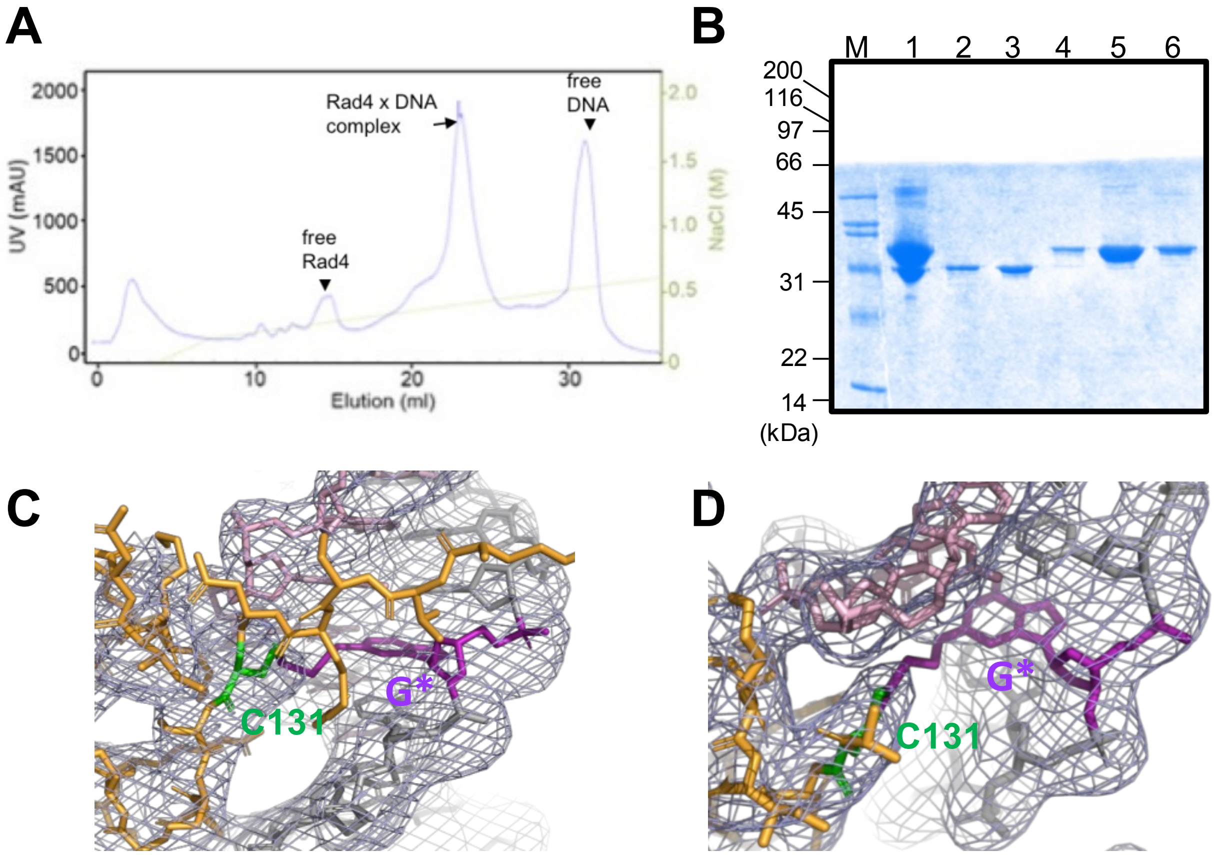

**Figure S1. Purification and crystallization of Rad4 mutant proteins tethered to CCC/GGG DNA. (A)** The mutant Rad4 (Δβ–hairpin3)–Rad23 crosslinked to CCC/GGG_G* DNA was purified over a MonoQ column (GE Healthcare) over a 0–2 M NaCl gradient. The peaks corresponding to free Rad4-Rad23 ('free Rad4'), crosslinked complex ('Rad4 x DNA'), and free DNA are indicated with arrows. **(B)** Non-reducing SDS-PAGE gel of the eluted fractions shows that the crosslinked Rad4-Rad23-DNA complex (Lanes 4-6; eluting at 400–480 mM NaCl) was separated from the free Rad4-Rad23 (Lane 2, 3; eluting at 280–320 mM NaCl). Lane 1 shows the input. **(C, D)** 2Fo-Fc map of the region near the crosslink at a contour level of 2σ for Δβ–hairpin3 x CCC/GGG complex (C) and for Δβ–hairpin2 x CCC/GGG complex (D).

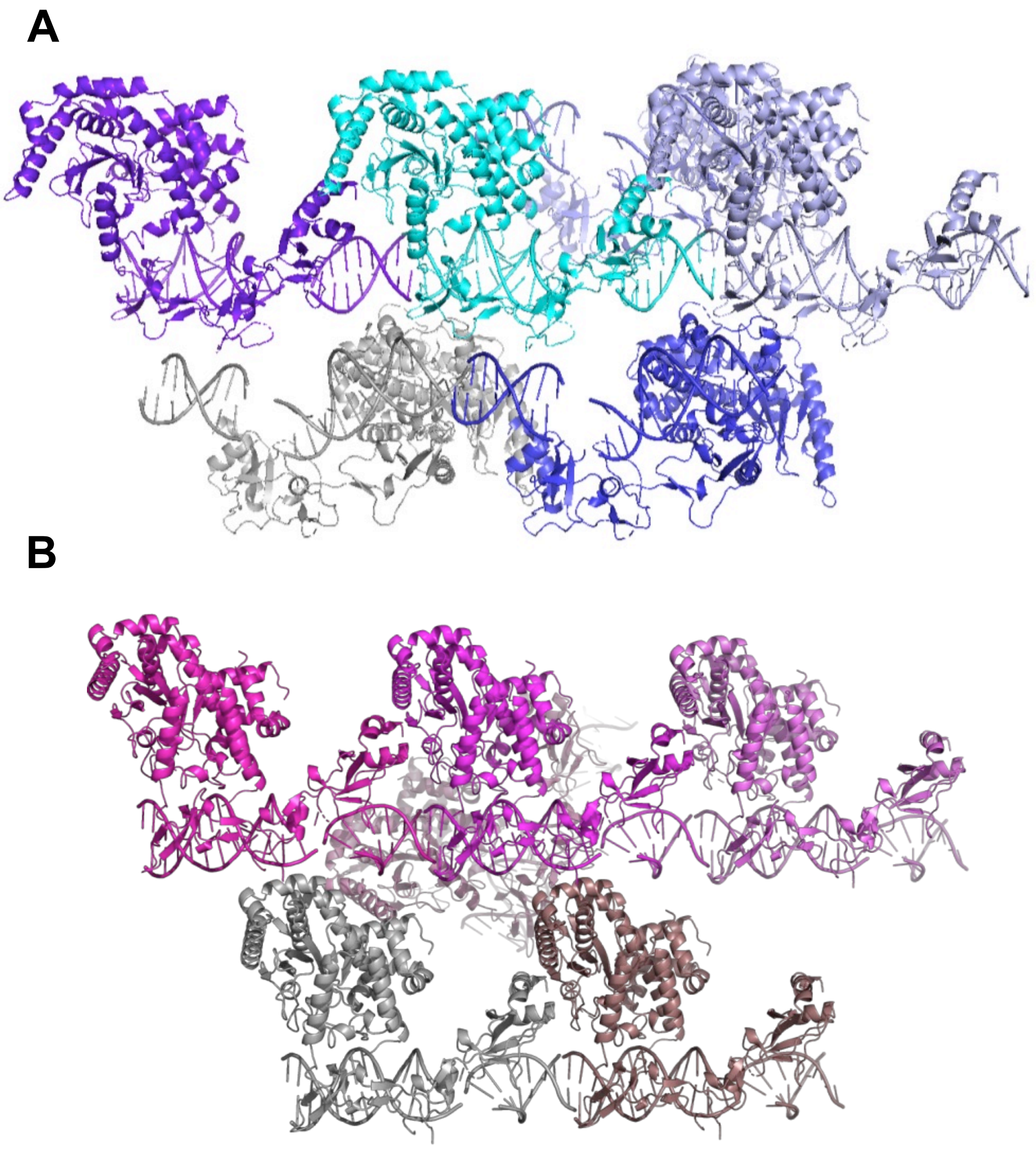

**Figure S2. Crystal packing arrangements of the Δβ−hairpin3 and Δβ−hairpin2 tethered to CCC/GGG DNA.** The head-to-tail arrangements of molecules in the Δβ−hairpin3-DNA **(A)** and the Δβ−hairpin2-DNA crystals **(B).** The symmetry-related, neighboring molecules show the differences in the packing arrangements between the two crystals.

A

B

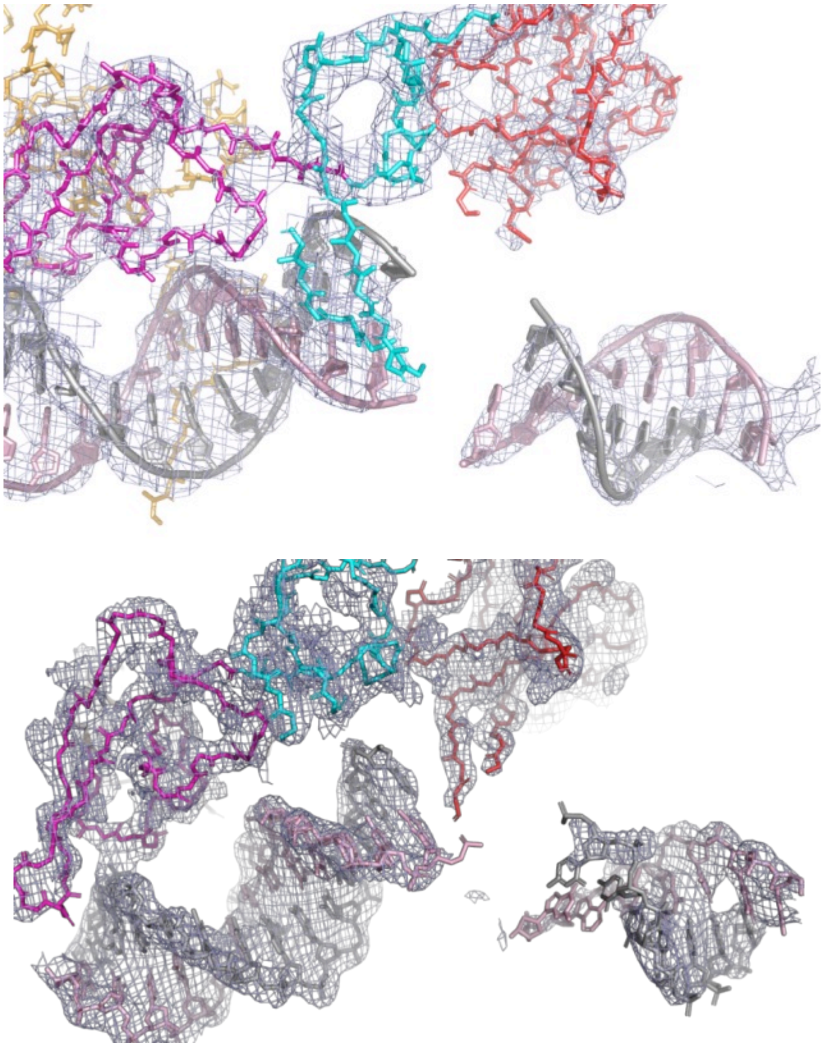

**Figure S3. Electron density showing the 'open-like' conformation of DNA.** 2Fo-Fc electron density maps showing the 'open-like' DNA conformations in the Δβ−hairpin3-DNA complex **(A)** and in the Δβ−hairpin2-DNA complex **(B)**. Maps are shown at a contour level of 1.5σ.

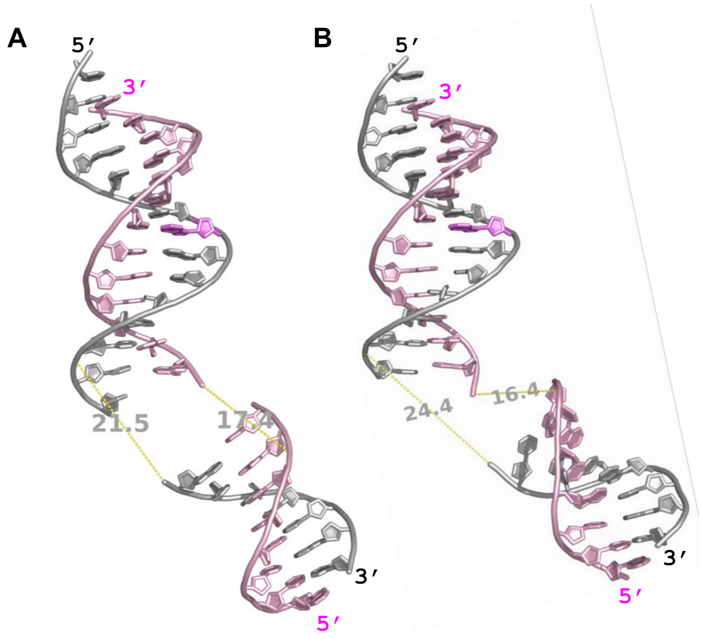

**Figure S4. DNA in the 'open-like' structures.** **(A)** Δβ–hairpin3 and **(B)** Δβ–hairpin2. The distances between the backbone phosphorus atoms of the common visible nucleotides are shown for top strand (W14 and W18) and the bottom strand (Y7 and Y12) in both structures.

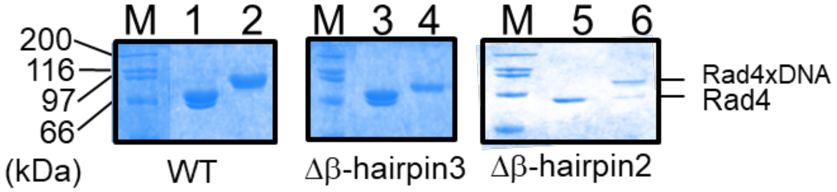

**Figure S5. Non-reducing SDS-PAGE for the crosslinked complex used in FLT measurement.** Upon crosslinking, free nontethered DNA was removed from the tethered complexes either by MonoQ column or extensive dialysis (MWCO 50 kDa) in high salt (800 mM NaCl) conditions. Samples were then analyzed on 15% SDS-PAGE gels using a loading buffer lacking 2-mercaptoethanol. Lane M indicates molecular weight marker. Lanes 1 & 2 show samples with WT Rad4 without and with tethered the CCC/GGG_G*_DA duplex, respectively. Lanes 3 & 4 are analogous samples with Δβ–hairpin3, and Lanes 5 & 6 are with Δβ–hairpin2. The tethered Δβ–hairpin2 samples contained some excess nontethered proteins. However, such nontethered protein does not impact FLT even if it interacted nonspecifically to the CCC/GGG_G* DNA (see **Figure 4**).

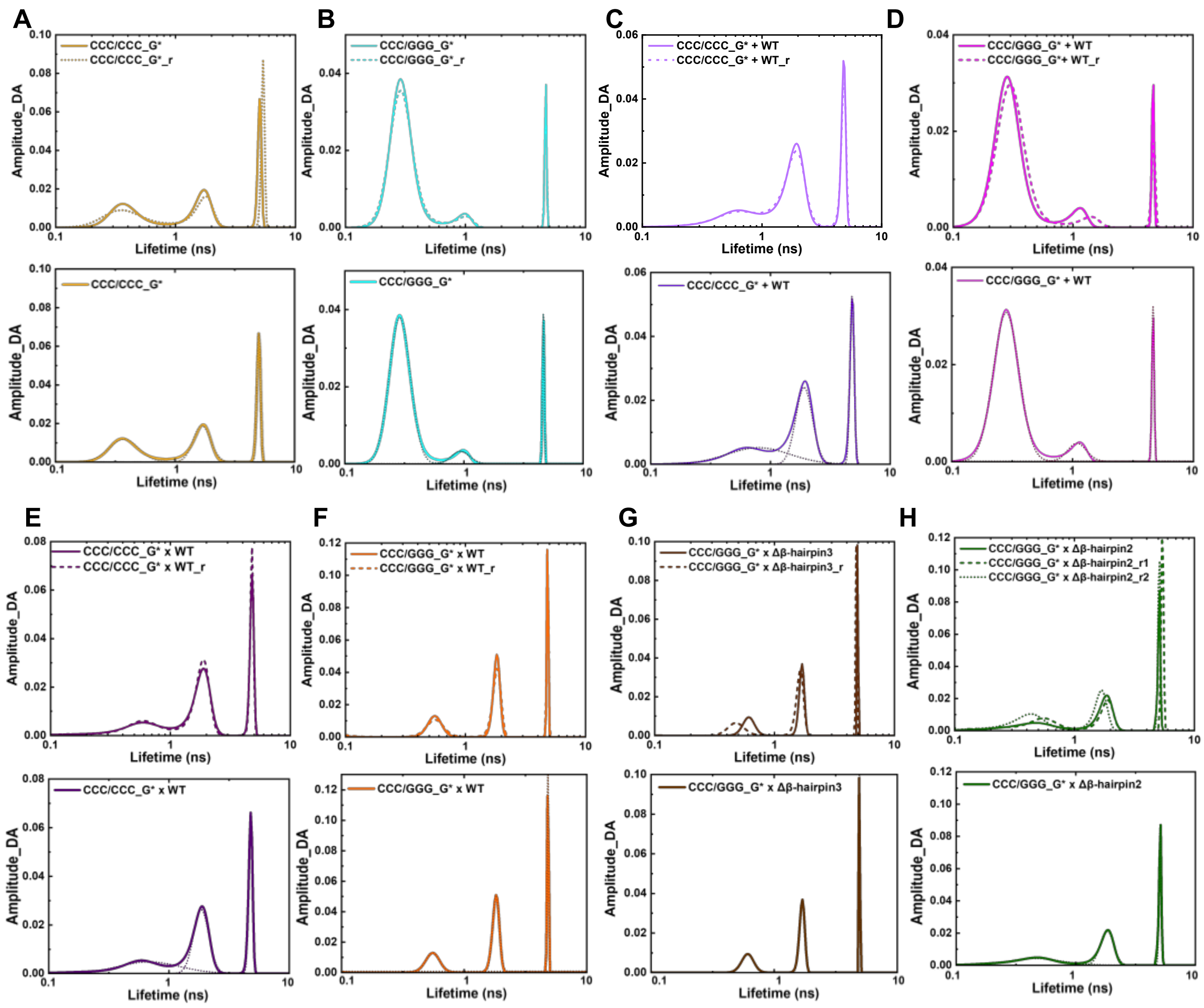

**Figure S6. Fluorescence lifetime distributions by MEM analyses and Gaussian fitting.** Top panels show the reproducibility of the fluorescence lifetime distributions obtained by MEM. Solid and dotted lines show different replicates of each sample. Bottom panels show the Gaussian-fitted peaks (black dotted line) underlying the representative profile (solid line). **(A)** Mismatch CCC/CCC_G* (yellow); **(B)** matched DNA, CCC/GGG_G* (cyan), **(C)** CCC/CCC_G* noncovalently bound to WT Rad4 (violet), **(D)** CCC/GGG_G* noncovalently bound to WT Rad4 (magenta), **(E)** CCC/CCC_G* covalently tethered to WT Rad4 (deep purple), **(F-H)** CCC/GGG_G* tethered to WT Rad4 (F; orange), to Δβ−hairpin3 (G; brown), or to Δβ−hairpin2 (H; green). All amplitudes indicate the normalized, fractional amplitudes. Full reports of the lifetimes, fractional amplitudes, FRET efficiencies of each Gaussian peak from all replicates as well as the sample's average FRET efficiencies are in **Table S2**.

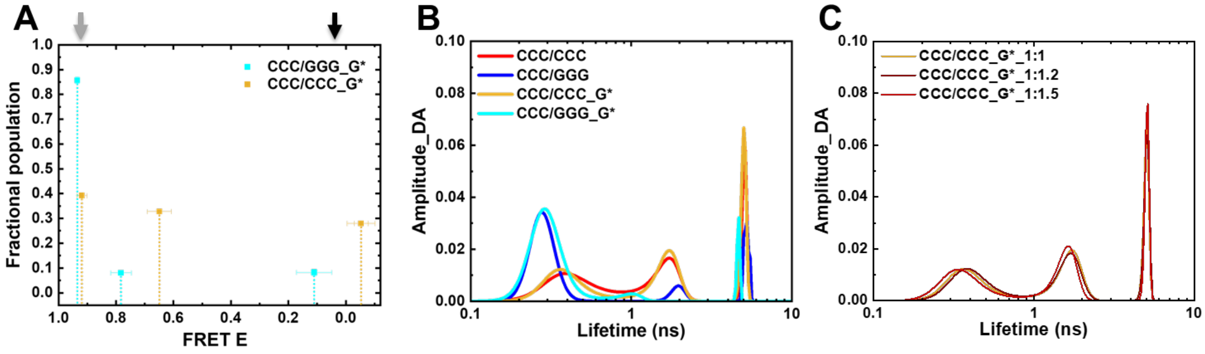

**Figure S7. FRET and fluorescence lifetime (FLT) distributions of the tC$^{o}$-tC$_{nitro}$-labeled DNA. (A)** The FRET efficiencies and its fractional populations obtained from MEM-Gaussian fitting for CCC/CCC_G* (yellow) and CCC/GGG_G* (cyan). The arrows indicate the computed values of FRET for B-DNA conformation (gray) and for the DNA conformation in the Rad4-bound ‘open’ crystal structure (black), computed as described in Methods. CCC/GGG_G* adopts mostly B-DNA conformation whereas CCC/CCC_G* shows heterogeneous conformations with three major peaks: high FRET, medium FRET and low/zero FRET. Low/zero FRET portion overlaps with the FRET expected of the ‘open’ conformation in crystal structure. Horizontal and vertical error bars indicate s.d of the values from 2 independent measurements. **(B)** FLT distributions for donor/acceptor-labeled DNA: CCC/CCC without G* (red), CCC/CCC_G* (yellow), CCC/GGG without G* (blue) and CCC/GGG_G* (cyan). Similar profiles between DNA without and with G* indicate that the presence of G* does not significantly alter the DNA conformations as sensed by the current probe position. All amplitudes indicate the normalized, fractional amplitudes. **(C)** The FLT distributions of CCC/CCC_G* DNA annealed with varying donor:acceptor strand ratios. Donor and acceptor strands are the bottom and top strands in Figure 4A, respectively. The FLT distributions remain the same among these samples indicating that the low/zero FRET peak is not due to an excess, unannealed donor strand in the sample.

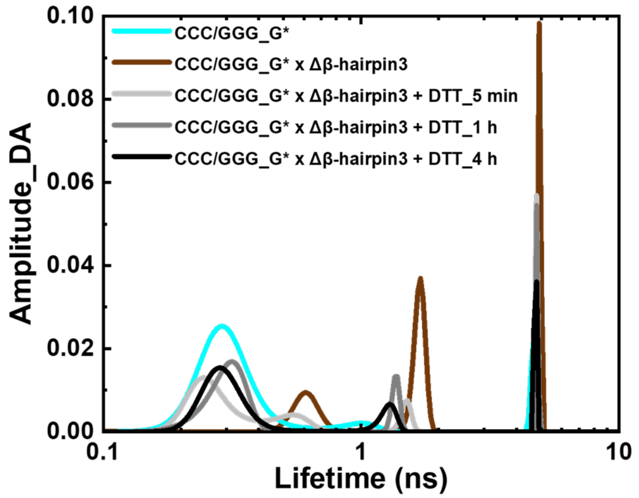

**Figure S8. DTT treatment showing the reversibility of crosslinked complex formation.** The MEM lifetime distribution of the Δβ–hairpin-3 x CCC/GGG_G* complex treated with DTT shows that the tethering is reversible. The crosslinked complex (brown) was treated with 5 mM DTT and FLT measured after 5 mins (light grey), 1 h (grey) and 4 h (black) respectively. The change in conformational distribution after DTT treatment supports the reversibility of the 'open-like' complex in the presence of DTT.

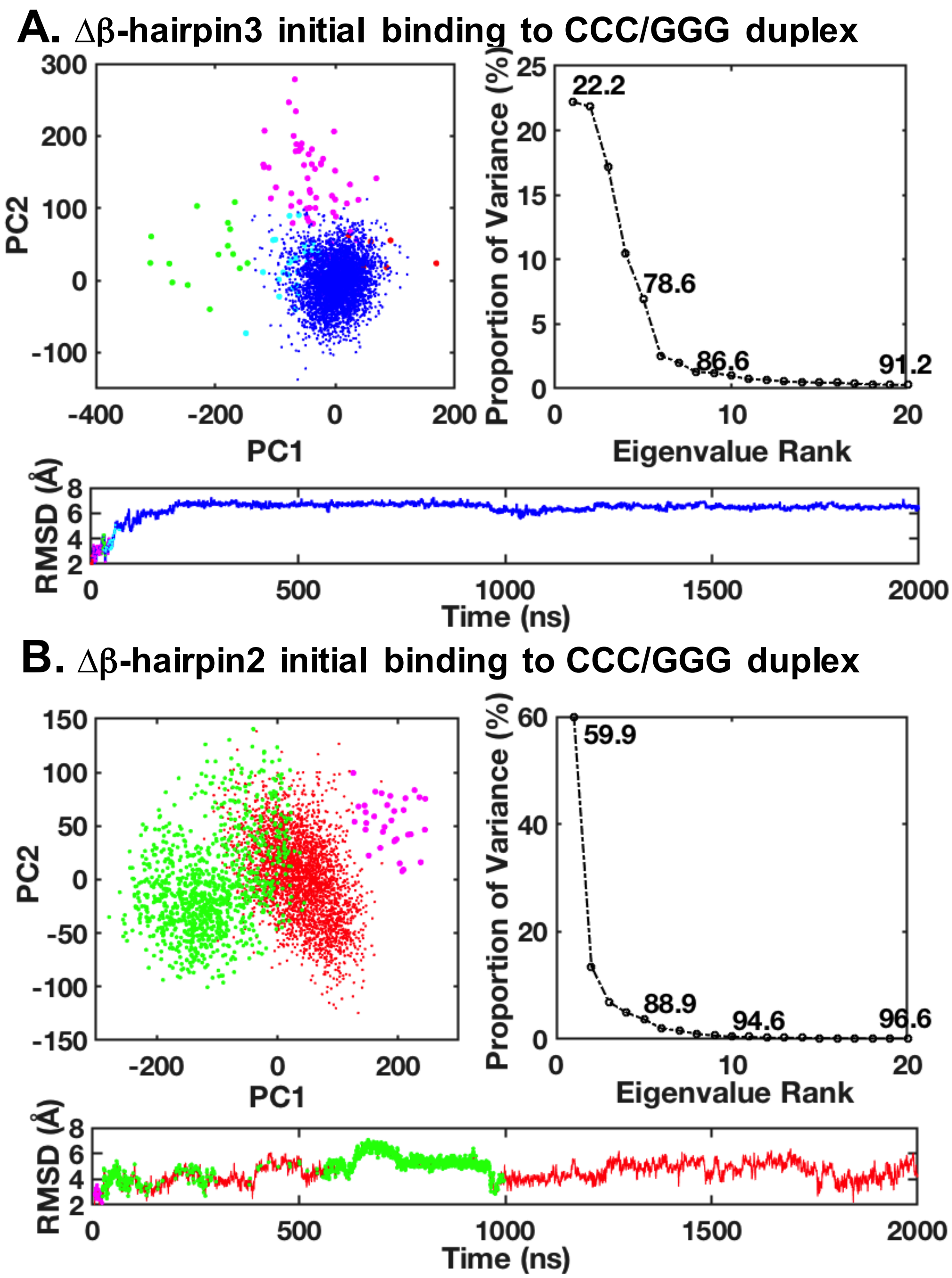

**Figure S9. Principal component analyses and RMSD for the MD simulations of the Rad4 mutants initial binding to the CCC/GGG duplex. (A)** Δβ−hairpin3 and **(B)** Δβ−hairpin2. The 1 to 2 μs ensemble was used for further analyses since one main structure cluster is achieved in ~ 1 μs (with stable RMSD value). Different structure clusters are color-coded. The RMSD values of best representative structures for the ensembles between 1-2 μs are shown in black circles.

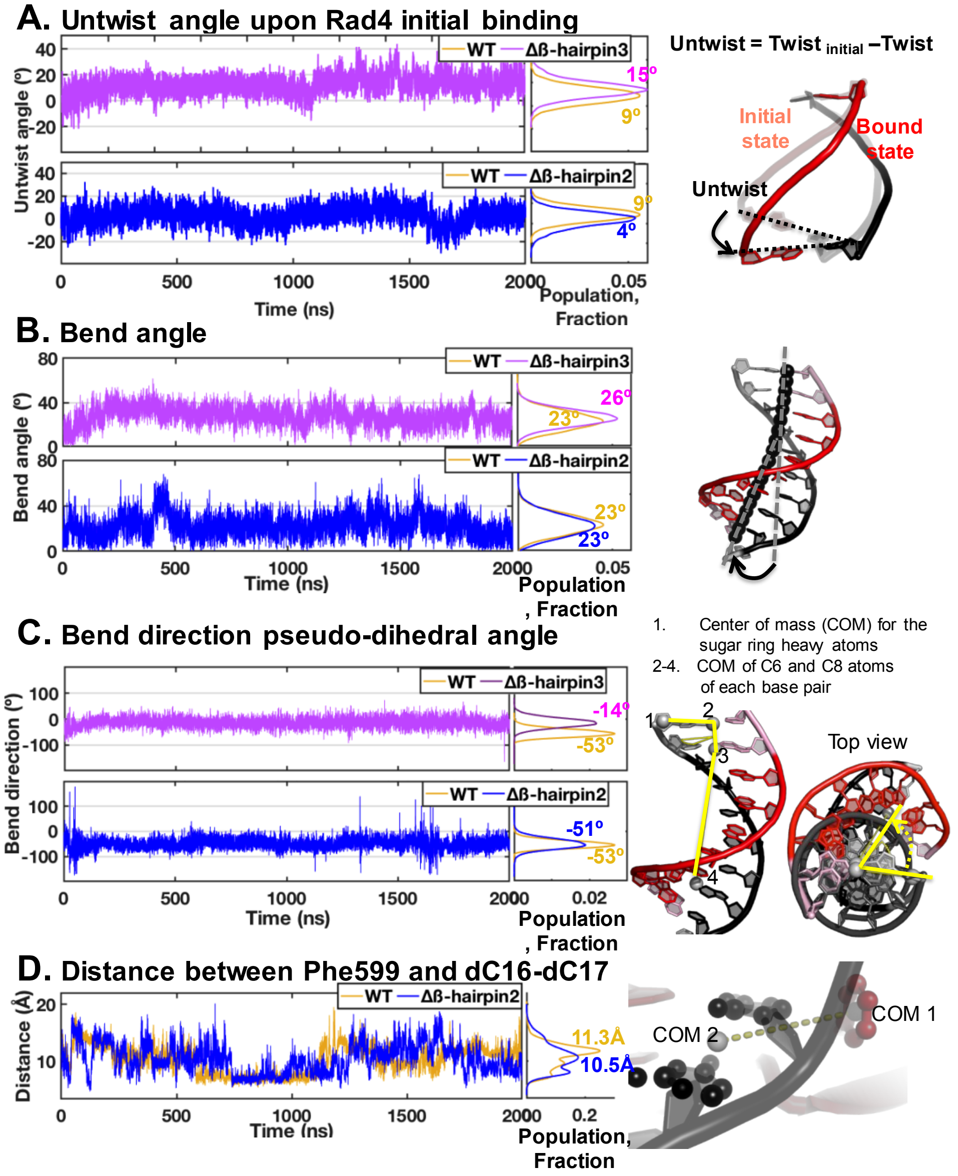

**Figure S10. DNA untwisting and bending analyses for the MD simulations for initial binding of Rad4 mutants to the CCC/GGG duplex.** **(A)** DNA untwist angles, **(B)** bend angles, **(C)** bend directions, and **(D)** distances between Phe599 and the partner bases at the potential 'open' site (see SI Methods for the definitions). Time-dependent trajectories (left) and kernel densities (right) are shown for all properties. Kernel densities for the values of the initial binding states (1-2 µs) are calculated using the ksdensity function with 200 bins in MATLAB 7.10.0 (The MathWorks, Inc.), and are plotted on the side with mean values labeled for (A-C) and median values labeled for (D). These kernel densities are representative of the population distributions over the range of each property (e.g. untwist angle).

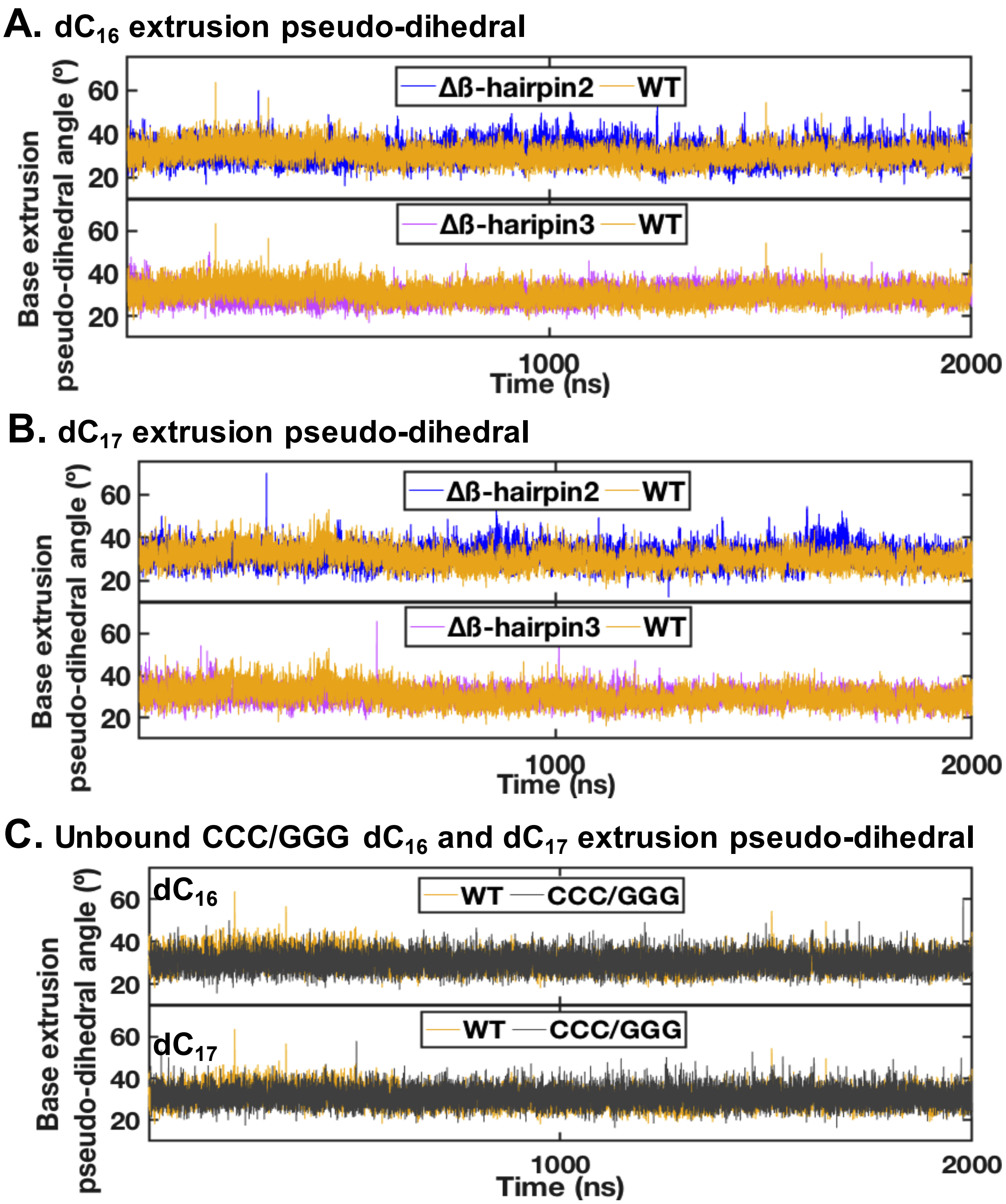

**Figure S11. Partner strand cytosine base extrusions in different Rad4-DNA complexes. (A)** $dC_{16}$ and **(B)** $dC_{17}$ extrusion pseudo-dihedral angles show that only Δβ–hairpin2 promotes the partner base extrusions more than in the WT or (**C**) the unbound CCC/GGG DNA (also see **Figure 4**). The values for the WT and unbound DNA are computed from the trajectories in ref. (17).

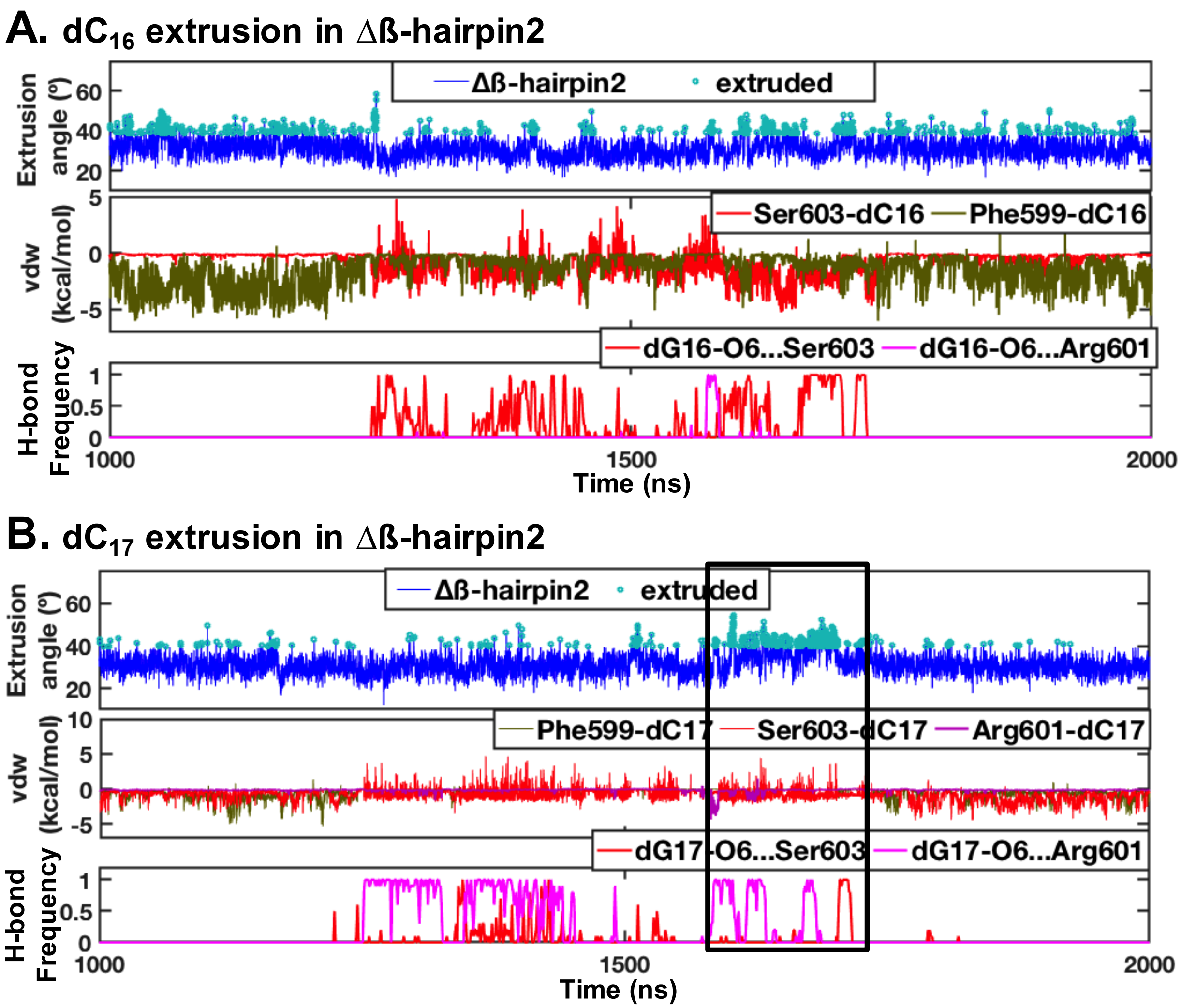

**Figure S12. Structural and energetic origins of enhanced DNA base extrusions in the Δβ−hairpin2-DNA complex.** The extrusions of $dC_{16}$ **(A)** and $dC_{17}$ **(B)** in the partner strand (W16 and W17, Figure 2A) are revealed in the base extrusion pseudo-dihedral angles. The extrusion of $dC_{16}$ correlates with the van der Waals (vdw) interaction energies between Phe599 and $dC_{16}$. The extrusion of $dC_{17}$ correlates with the frequency (fraction of ensemble with hydrogen bonds/ns) of hydrogen bonding between β−hairpin3 tip residues, Arg601 and Ser603, and the guanine opposite $dC_{17}$ (termed '$dG_{17}$'); the high frequency hydrogen bonding weakens the Watson-Crick hydrogen bonding between the dG-dC base pair and hence facilitates $dC_{17}$ extrusion. The boxed region highlights the correlation.

**SI Movies S1-S2**

**Movie S1.** Best representative structure for CCC/GGG DNA upon initial binding with Δβ -hairpin3, showing the greater engagement of BHD2 with the DNA minor groove compared to WT-Rad4. (Movie S1_b-hairpin3 mutant_walkaround.mp4)

**Movie S2.** Best representative structure for CCC/GGG DNA upon initial binding with Δβ-hairpin2, showing limited BHD2 hairpin insertion in the minor groove but close interaction of intact BHD3 hairpin with the major groove. (Movie S2_b-hairpin2 mutant_walkaround.mp4)
